# Mapping the Competence Boundary of a Protein Property Model: A Case Study on Plastic-Degrading Enzymes using ProtTrust-XAI

**DOI:** 10.64898/2026.08.01.742252

**Authors:** Abdulmujeeb T. Onawole, Rukayat O. Adegoke

**Affiliations:** Institute for Molecular Bioscience, The University of Queensland, Brisbane, 4067 Australia; Department of Pure and Applied Biology, Ladoke Akintola University of Technology, PMB 4000, Ogbomoso, Nigeria

**Keywords:** Protein property prediction, Prediction reliability, Protein model, Explainable AI, Applicability domain

## Abstract

Machine learning models for protein properties are usually reported by a single accuracy figure, which says how a model behaves on average but not whether to act on any one prediction, especially for a protein unlike anything in the training set. That gap is both a black box problem and an out-of-distribution problem, and it is worst exactly where discovery work happens, on sequences the model has not seen. We present ProtTrust-XAI, a framework that scores each prediction by ensemble consensus and by the structural coherence of its own attribution, and separately tracks a third signal, distance from the training distribution, to catch cases the first two cannot see. We demonstrate it on per-protein thermostability, training a relational graph convolutional network on melting temperatures for over 20,000 proteins using AlphaFold-derived contact graphs and frozen protein language model embeddings. On family-level held-out proteins the model reaches a Spearman correlation of 0.65 and a mean absolute error of 4.1°C, and predictions the framework labels most trustworthy fall to 3.0°C, below the assay’s own reproducibility floor, so a practitioner can act on the label with the same confidence as on the measurement itself. Applying the framework across the full dataset also exposes two representational blind spots, one around cofactor chemistry and one around membrane proteins, each with a distinct mechanistic explanation that points to a specific fix. Transferred to plastic-degrading enzymes at low sequence identity to the training data, absolute predictions collapse while the ranking survives, and a controlled ablation shows this is a general property of distribution shift rather than something particular to that external set. The same transfer identifies where the distance-based signal itself needs recalibrating before deployment, which is a diagnosis the framework produces about itself and not a hidden failure. The result is a practical rule. Inside a model’s competence domain, trust its labels. Outside it, trust its ranking. A model that reports its own limits, rather than only its average accuracy, is one an experimentalist can actually build on.

## 1. Introduction

Protein property prediction has been transformed by pretrained sequence models [1] and by accurate structure prediction [2], and models are now routinely reported by a single average accuracy across a benchmark. That number answers a question a practitioner rarely asks. The decision facing a structural biologist or a protein engineer is not how a model performs on average but whether to act on one specific prediction, for one specific protein, often a protein unlike anything in the training set. A point estimate carries no signal about that, and the gap is worst precisely where the stakes are highest, since discovery campaigns are by definition run on unfamiliar sequences. This work asks where the boundary of a model’s competence lies, and how that boundary can be measured rather than assumed. The same absence of a decision-usable signal has stalled explainable AI in an adjacent setting, drug discovery, where explanation methods have lacked standardised evaluation criteria and so have not been trusted enough for routine use in hit optimisation [3]. The same question, knowing when a prediction can be trusted, applies with particular force to protein engineering for plastic degradation, the case study this paper builds toward.

Global plastic production runs into the hundreds of millions of tonnes a year, and enzymatic recycling is one of the few routes that can break this material down without the energy cost of incineration or the quality loss of mechanical recycling [4]. Polyethylene terephthalate (PET) hydrolases are the enzymes that do this work, cleaving the ester bonds that hold polyethylene terephthalate together, and their industrial usefulness depends on surviving the elevated temperatures that soften PET enough for the enzyme to reach it. Thermostability is therefore not incidental to this application, it is the property that decides which candidate enzymes are worth expressing and testing, which is why AI-assisted discovery for plastic-degrading enzymes has become an active research direction in its own right [5]. Knowing when a thermostability prediction can be trusted is therefore not an abstract question about model evaluation, it decides which enzymes get sent to the bench. Answering it needs a property for which the question can actually be settled, and per-protein thermostability is an unusually demanding testbed in its own right, since uniform measurements exist at scale, distribution shift can be quantified rather than asserted, and the property itself is a global emergent feature of an entire fold rather than a locally determined one, which makes it a demanding case for per-residue attribution and, as we show later, exposes an explanation method that does not survive the move from small molecules to proteins. Plastic-degrading enzymes, introduced above as the reason thermostability matters, become the case study later in the paper through which we stress-test the competence boundary this framework measures.

The framework itself is not specific to thermostability. Its three axes are defined on ensemble behaviour, attribution geometry and embedding distance, none of which reference the predicted quantity, so the same construction applies to solubility, expression, binding or any protein property where an ensemble and a per-residue attribution can be computed. The demonstration here is melting temperature. The object of study is the reliability of protein property models under shift. We previously built a two-axis version of this idea, ensemble consensus and the coherence of a model’s own attribution, and validated it on the electronic properties of transition-metal complexes [6]. Two axes are not enough here, for a known reason. Ensemble members trained on partitions of the same data can share a bias, so their agreement measures how consistently a model reaches an answer rather than whether that answer is right [7, 8]. Under mild shift the two axes stay correlated enough for consensus to be a useful proxy, but far from the training distribution they can come apart entirely. This work adds a third axis measuring distance from the training distribution, and tests all three in a setting where the shift is large and quantifiable rather than assumed. This gives four contributions. We train a structure-grounded model in which every protein carries its own contact graph, rather than the single fixed backbone shared across variants that a mutation-effect model typically uses, which makes architectural choices consequential and lets them be searched rather than fixed by assumption. While recent structure-grounded architectures such as DeepFRI [9] and SSRGNet [10] demonstrate the power of combining protein language models with graph neural networks for function and secondary structure prediction, they rely strictly on point-estimates and aggregate benchmark accuracy. ProtTrust-XAI extends this lineage by adding a calibrated, 3-axis reliability taxonomy on top of a relational structural encoder. Rather than using explainability merely to visualize active residues post-hoc (as in DeepFRI), we repurpose attribution geometry as an active diagnostic instrument that quantifies the model’s competence boundary under severe distribution shift. We add a third, applicability-domain axis so that agreement between ensemble members is checked against distance from the training distribution, rather than trusted on its own. The resulting trust taxonomy is built to function as a diagnostic instrument, one capable of exposing where a model’s representation is thin rather than only how accurate it is on average. A competence boundary is only meaningful if it can be tested against data the model has never seen. We measure that boundary directly against an external enzyme set and a taxonomy-only ablation, so that what survives real distributional shift, and what does not, is shown rather than assumed. Industrial enzyme engineering follows from this boundary as one application of it, not as the reason it was built. The value of a framework like this is not a marginal gain in accuracy over the next model. It is speed of decision. A practitioner who knows which predictions to act on, which candidates to test and which to set aside can move directly from a ranked list to a bench, without spending assay budget on a wrong answer along the way. That is the more useful currency, and giving a model the means to earn it is what this work sets out to do.

## 2. Methods

### 2.1 Data Curation

Training data were per-protein melting temperatures from Meltome Atlas [11] accessed through the FLIP benchmark [12], comprising 27,951 proteins with a measured melting temperature spanning 25.3 to 99.0°C. Proteins longer than 1,024 residues were excluded, and each remaining protein was matched to its AlphaFold model [13], requiring an exact agreement between the structure length and the sequence length, which yielded 24,194 per-protein graphs. Because the embedding cache was keyed on a hash of the sequence, 4,011 exact duplicate sequences collapsed to a single entry, leaving 20,183 unique proteins. This filtering and deduplication path, along with the external test set introduced below, is summarised in Table 1.

**Table 1.** Dataset composition and the external test set.

| Item | Quantity | Value |
| --- | --- | --- |
| Training pool (Meltome via FLIP) | Proteins with T<sub>m</sub> | 27,951 |
| After length ≤1024 + AlphaFold structure | Per-protein graphs | 24,194 |
| After exact-sequence deduplication | Unique proteins | 20,183 |
| Training pool | Proteins | 18,179 |
| Internal test (cluster-held-out) | Proteins | 2,004 |
| Internal test | Records (replicates) | 2,435 |
| Split basis | PIDE20 clusters, 20% identity | 14,185 |
| T<sub>m</sub> range (pool) | Degrees C | 25.3-99.0 |
| T<sub>m</sub> mean +/- std (pool) | Degrees C | 51.6 +/- 10.9 |
| Assay reproducibility floor | Median replicate spread, C | 3.89 |
| External PET set | Enzymes | 475 |
|  | With measured T<sub>m</sub> | 233 |
|  | Fully endpoint-labelled | 190 |
|  | Industrial hits | 26 |
| PET vs Meltome identity | Median global identity | 10% |
| PET structures | Esmfold, folded/attempted | 475/475 |
|  | Mean plddt* | 87.9 |

The 20,183 unique proteins were split into a training pool of 18,179 and an internal held-out test set of 2,004 proteins across 2,435 records, held out at the level of sequence-identity clusters rather than individual proteins. The external set comprised 475 PET-hydrolysing enzymes with measured expression, thermostability and depolymerisation activity, discovered across three screening rounds of 94, 191 and 190 candidates [14]. Of these, 233 carried a measured melting temperature and 190 carried all three endpoints. These enzymes sat at a median 10% global sequence identity to the training data, with 98% below 25% identity, and the training data contained no cutinases or PETases, so the external set was novel at the level of the protein family while remaining familiar at the level of the fold through 267 esterases and 101 lipases. Figure 1 compares how the training pool and the external PET set line up on melting temperature and sequence length.

**Figure 1.**
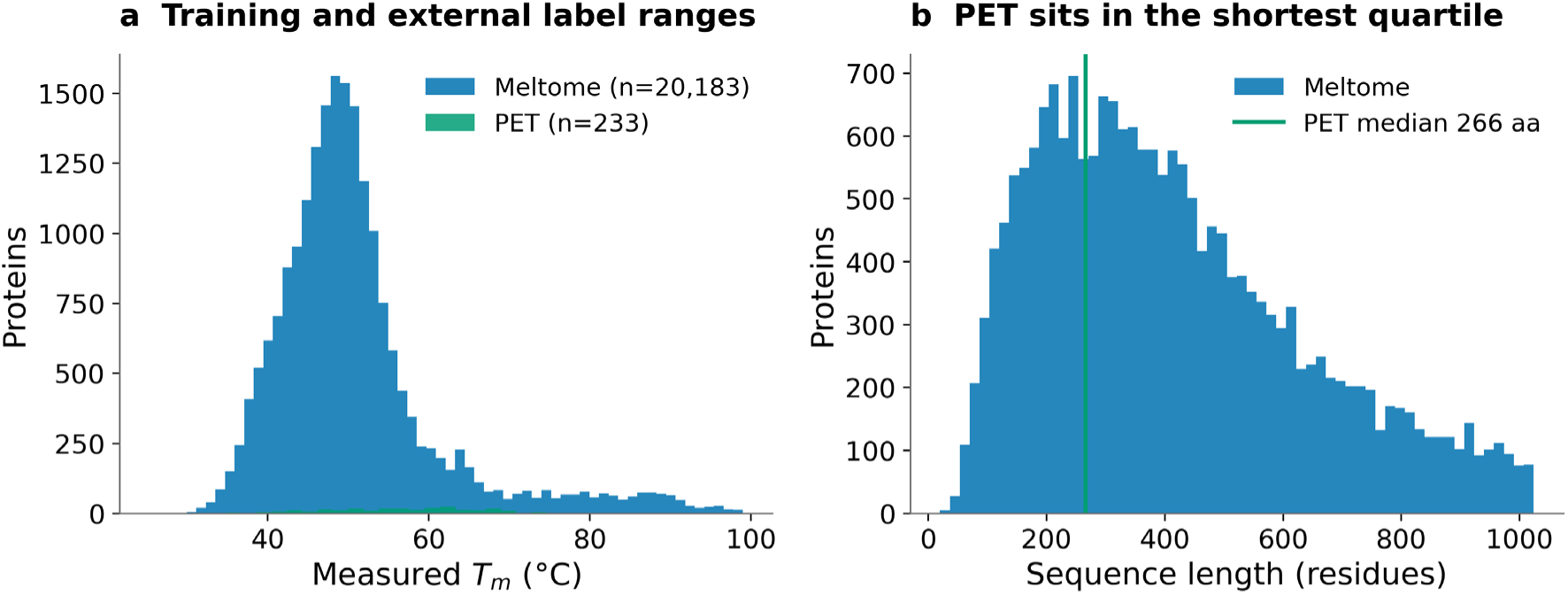
Dataset composition and the split. (a) Measured melting temperatures for the training data and the external test set. (b) Sequence-length distribution; the external set sits in the shortest training quartile.

Splits were built on FLIP’s pre-computed clusters at 20% sequence identity, so held-out proteins were novel at the level of the family rather than close homologues. This choice mattered because a random split on the same data permitted training and test proteins up to 94% identical, so an accuracy computed that way would largely describe interpolation among relatives rather than generalisation to a genuinely new protein. Every internal performance figure reported later in this paper is measured on this family-level split unless stated otherwise. Those duplicates were informative rather than merely redundant. Meltome measured the same sequence in multiple lysates and cell lines, and among 2,580 duplicate-sequence groups not one reported an identical melting temperature. The median spread within a group was 3.89°C, which we take throughout as the assay’s own reproducibility floor and as the irreducible error against which any reported error should be read. All 2,580 groups fell inside a single identity cluster, so the cluster split already absorbed them and no exact duplicate crossed a train and test boundary.

### 2.2 Model and training

Each protein was represented as a graph whose nodes are residues carrying frozen 1,280 dimensional ESM2-650M embeddings [15]. Two edge relations were defined, a backbone relation for sequential connectivity and a contact relation for spatial proximity, with a contact edge placed between residues whose C-β atoms lie within 8.0 Å, measured at C-α for glycine, following established convention [16]. Separating covalent connectivity from fold-induced proximity allowed the relational network to learn distinct weights for the two interaction types. The model is a relational graph convolutional network [17] with a single regression head predicting standardised melting temperature. Architecture and optimiser settings were selected by 40 Optuna trials [18] on a family-level cluster fold, pre-registered before cross-validation, searching learning rate, weight decay, batch size, number of relational layers, node dimension and dropout. The search selected two relational layers, a node dimension of 128, a learning rate of 3.51×10⁻⁴, a weight decay of 5.67×10⁻³, a batch size of 16 and a dropout of 0.051 (Table S2).

The depth result was a clean negative rather than a null one. Two layers won, and every six-layer trial sat at the bottom of the table, a measured over-smoothing signal of the kind reported for naive deep graph networks [19, 20] and only partly mitigated by the residual connections our blocks carried. This task wanted a shallow residue graph. Early stopping rather than dropout was the effective capacity control, since the best epoch fell between 4 and 8 in every one of the 15 training runs. Melting temperatures were standardised using statistics fitted on the training pool alone. Model quality was estimated by 5-fold, 3-repeat cross-validation at the level of identity clusters, and the deployed predictor was an ensemble of five models, one per fold, selected as the best checkpoint across repeats.

Hyperparameter search used one cluster fold of the training pool as its validation partition. That partition coincided with the first cross-validation fold, because both were drawn with the same seed from the same pool, so the corresponding checkpoint’s cross-validation score carried the optimistic bias that arises when tuning and evaluation share a partition [21, 22]. The held-out test set was withheld before the pool was formed and never entered the search, so no reported test metric was affected. The overlap’s own scale bounded its consequence: the affected fold scored 0.6714 in cross-validation, within 0.0007 of two other folds with no such overlap, and excluding it moved the cross-validation mean from 0.6554 to 0.6542, well inside the fold-to-fold standard deviation of 0.0122. Across all fifteen checkpoints, cross-validation score and held-out test performance showed no resolvable relationship (Spearman rho −0.15, bootstrap 95% CI −0.67 to 0.35), so no fold’s cross-validation rank, including the affected one, reliably predicted its held-out rank. The affected checkpoint ranked 13th of 15 on the held-out test set despite ranking first in cross-validation, the opposite of what a meaningfully advantaged checkpoint would show. The five-member ensemble outperformed even its single best member (0.6476 versus 0.6435 Spearman), indicating that averaging across checkpoints, rather than which specific checkpoints were chosen, is what the ensemble gained from.

Five models were used primarily for practical reasons. Computing occlusion attributions, described later, is expensive per model, and five was a reasonable ensemble compute without becoming impractical. The number also fell out naturally from the 5-fold cross-validation already used for model selection, one model per fold, so no additional models needed to be trained beyond what cross-validation already required. This choice turned out to serve a second purpose beyond cost. Since the five members are separately trained rather than pooled, their agreement on a given protein is itself a signal the consensus axis draws on later, which a single pooled model cannot provide.

### 2.3 Protein-weighted evaluation

Meltome contained multiple measurement rows for many proteins, with a maximum of 11 replicate rows for a single protein. Because replicate rows shared a sequence and therefore a graph, the model produced an identical prediction for all of them while their measured values differed, so scoring at the level of records both up-weighted proteins with more replicates and charged the model the assay’s own irreducible noise, since a single prediction could not match several differing measurements at once. All primary metrics therefore averaged replicate labels to one value per protein instead, giving 2,004 test proteins from the original 2,435 records, with record-weighted figures reported in Supporting Information. A control confirmed this was justified. Averaging replicate labels also reduced label noise and could therefore be accused of flattering the model, so a single randomly sampled row per protein was scored instead, which did not reduce label noise. That control recovered a Spearman correlation of 0.6445 against the averaged 0.6476, so only a small part of the gain over record weighting was noise averaging, and the remainder came from removing duplicate weighting. Deduplication was applied to evaluation only; training retained replicates as documented sample weighting.

### 2.4 Reliability axes and explanation attributions

Every prediction carries a label from two crossed binary axes, giving the four scenarios A, B, C and D used in our earlier work [6]. Consensus is high when the ensemble members agree closely on the prediction. Coherence is high when the attribution concentrates on the structurally important residues, meaning the buried, densely packed core. Scenario A is high consensus and high coherence, B is high consensus alone, C is high coherence alone, and D is neither. The consensus axis is the standard deviation of the five ensemble predictions divided by the standard deviation of predictions across the calibration cohort, a form adopted because the target was standardised and centred near zero, where a conventional coefficient of variation diverges. A prediction is defined as high consensus when this ratio falls below the 30th percentile of the calibration cohort.

The coherence axis measures whether attribution mass concentrates on the buried, densely packed core, an established structural correlate of thermostability [23, 24]. A whole-protein property has no mutated site on which to centre a local explanation, so the analogue adopted here was the localisation of attribution magnitude over the residues of highest contact number. A prediction is defined as high coherence when this localisation exceeds the 70th percentile of the calibration cohort. The core is defined as the fifth of residues with the highest contact number, and because contact numbers are small integers, several residues typically tie at that boundary, so which tied residues enter the core is resolved arbitrarily. We quantified the consequence by relabelling every protein under a different resolution of those ties. Individual coherence values moved by a mean of less than 0.001 and 3.1% of proteins changed scenario, while scenario coverage over the whole dataset moved by at most 0.04 % points. The tie resolution is therefore unbiased with respect to the reported distributions, though an individual protein’s label is reproducible only up to this choice. Its diagnostic value is tested directly later in this paper, where it proves more sensitive than the consensus axis to a missing structural representation.

The applicability-domain axis measures the distance from a query protein’s mean ESM2 embedding to its nearest training neighbour, with an empirical quantile cutoff, and exists to catch the case the consensus axis cannot see, in which every ensemble member is confidently wrong in the same direction because none of them has seen anything like the query [7]. All cutoffs are empirical percentiles of a calibration cohort drawn from the training pool, so a cutoff value has no meaning without the cohort that produced it. Each calibration therefore records a provenance block carrying the ensemble members, their checkpoint hashes, the label statistics, the split and graph constants and a hash of the analysis code, and downstream steps refuse to run against a calibration written by a different ensemble.

Per-residue attributions are computed by occlusion, setting a residue’s node features to zero and recording the change in prediction, following the substructure-masking approach [25]. Prediction uses all five ensemble members, but occlusion attribution, and therefore the coherence axis, runs on a single checkpoint, the member with the highest cross-validation score among the fifteen trained models. Because that choice could in principle shape the resulting taxonomy, its sensitivity was tested directly rather than assumed. The full labelling pipeline was recomputed once per attribution member, over the identical calibration cohort and the identical 2,004-protein test set (Supporting Information, Table S8). The ordering of error across scenarios replicated in every arm, with Scenario A carrying the lowest error throughout, and no protein moved between the trusted and untrusted band under any choice of attribution member, so the decision the framework supports was invariant to this choice. Agreement on the finer distinction within a band, a protein’s assignment to A rather than B, or to C rather than D, was weaker, at 69.7% to 75.6% depending on the pair, against a tie-order floor of 95.1% measured independently. Per-residue attributions agreed at a median rank correlation of about 0.20 between members, rising to 0.33 when disjoint subsets of members were averaged before comparison. Attributions are accordingly reported as population-level evidence rather than as a per-residue claim about any single protein.

That earlier framework sets four criteria for validating an explanation method before it is used downstream, and the coherence axis defined above is a construct with no ancestor in it, so it was tested rather than assumed. It was compared against the prediction-explanation direction criterion, adapted to regression in our transition-metal work, in which a prediction above the dataset mean should carry predominantly positive attributions [6]. Holding the consensus axis and all cutoffs fixed so that only the explanation axis changed, the two were compared on their ability to separate error, and the direction criterion was separately evaluated on its own terms as a per-protein consistency rate against a chance level of 0.50.

### 2.5 Baseline, ablation and external evaluation

Three further experiments test whether the framework holds outside the conditions it was built and calibrated on: a baseline comparison against a simpler model, a controlled ablation of distribution shift, and an evaluation on a wholly external protein set. A ridge regression on mean-pooled ESM2 embeddings was trained on the same pool with the same cluster splits and scored on the same proteins, with both trust axes computed on it identically, so that the framework, not just accuracy, is compared like for like. Because both models score the same proteins, the comparison is paired, resampling both models on a shared bootstrap index.

A leave-one-taxon-out ablation removed all 6,411 prokaryotes from training, together with any training protein sharing an identity cluster with the holdout, and retrained from scratch. A size-matched random cluster holdout was run as a control, because removing a taxon also removes training data and the two effects would otherwise be confounded. Both arms are ablations, and neither is the deployed ensemble. Structures for the external PET set are not in AlphaFold DB, and were predicted with ESMFold [15], yielding a structure for every candidate at a mean pLDDT of 87.9. Because the inherited cutoffs are raw values calibrated on a different distribution, the external evaluation reports two regimes. The strict regime inherits the cutoff values unchanged, and the rank-mapped regime applies the same percentile within the external cohort’s own distribution, which preserves the intended coverage when a distribution shifts in location or scale. Which regime will work can be diagnosed in advance, without labels, by comparing the empirical cumulative distributions of each axis before and after removing location and scale. A discrepancy that collapses under that standardisation indicates a shifted distribution of the same shape, where rank mapping restores the intended labelling. A discrepancy that survives indicates a genuine difference in shape that no rescaling repairs.

## 3. Results and Discussion

### 3.1 Evaluation metrics

The evaluation metrics on the internal performance showed that cross-validation gave a Spearman correlation of 0.6554 with a standard deviation of 0.0126 across the 15 runs. On the 2,004 held-out test proteins, which belong to identity clusters absent from every training fold, the ensemble reached a Spearman correlation of 0.6476, a root mean squared error of 5.48 °C and a mean absolute error of 4.10 °C (Figure 2).

**Figure 2.**
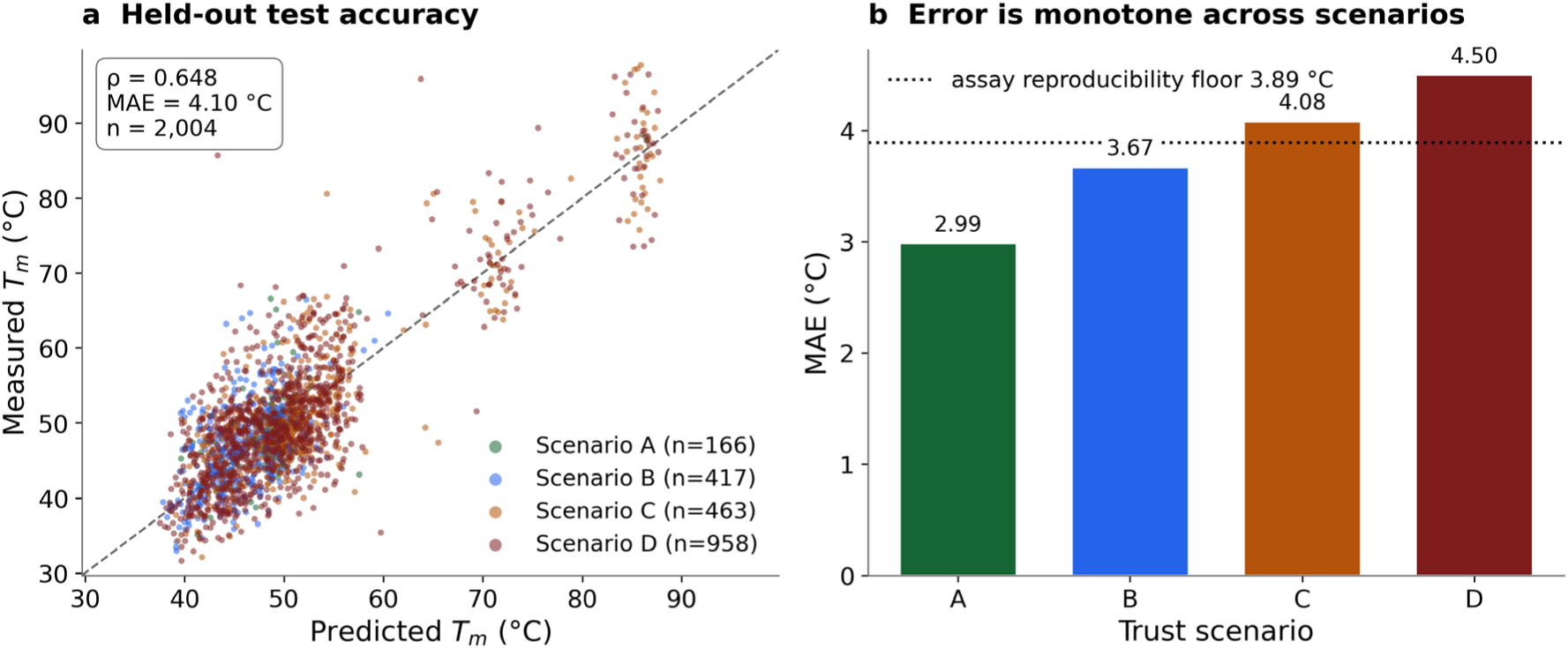
Held-out performance on the 2,004 test proteins, which belong to sequence-identity clusters absent from every training fold. (a) Predicted against measured melting temperature, coloured by trust scenario. (b) Mean absolute error by scenario against the 3.89 °C assay reproducibility floor measured from replicate assays of identical sequences.

The ensemble reached a Spearman correlation of 0.6476, higher than its best single fold (0.6435) or its mean single fold (0.6305). This gain did not come from selecting the best-performing members. Cross-validation rank showed no detectable relationship to held-out rank, with a Spearman correlation of −0.154 across the 15 checkpoints and only one of the top five by each criterion overlapping, consistent with no relationship at all at this sample size. A checkpoint’s cross-validation score therefore does not predict how well it generalises to a new protein family. The ensemble’s advantage comes from combining diverse members rather than from choosing good ones, which reinforces this paper’s broader point that in-distribution validation is a poor guide under shift.

The trust scenarios (Figure 2) show that error increases monotonically across the four scenarios, from a mean absolute error of 2.99 °C in Scenario A through 3.67°C and 4.08 to 4.50 °C in Scenario D, which is the ordering the framework predicts (Table 2). The trusted tier, Scenarios A and B combined, carries 3.47 °C against 4.36 °C for the flagged tier, a separation of 1.255 times. Consensus is the stronger of the two axes, separating error 1.256 times alone against coherence’s 1.119 times, and the two are near-independent, with a Spearman correlation between them of 0.05

**Table 2.** The consensus x coherence trust taxonomy on the 2,004 held-out test proteins, protein-weighted.

| scenario | n | Coverage (%) | MAE C | RMSE C | within scenario $\rho$ | measured Tm std C |
| --- | --- | --- | --- | --- | --- | --- |
| A | 166 | 8.28 | 2.99 | 4.07 | 0.398 | 6.24 |
| B | 417 | 20.81 | 3.67 | 4.86 | 0.523 | 5.65 |
| C | 463 | 23.1 | 4.08 | 5.39 | 0.707 | 11.86 |
| D | 958 | 47.8 | 4.5 | 5.97 | 0.664 | 10.76 |
| A+B | 583 | 29.09 | 3.47 | 4.65 | 0.509 | 5.87 |
| C+D | 1421 | 70.91 | 4.36 | 5.79 | 0.683 | 11.2 |
| ALL | 2004 | 100.0 | 4.1 | 5.48 | 0.648 | 10.15 |

Coverage and error answer only part of the practical question. A practitioner filtering candidates cares about enrichment (Figure 3), how much more often a trusted prediction actually falls within the assay’s own noise compared to chance, and that is not a quantity (Table 2) reports directly.

**Figure 3.**
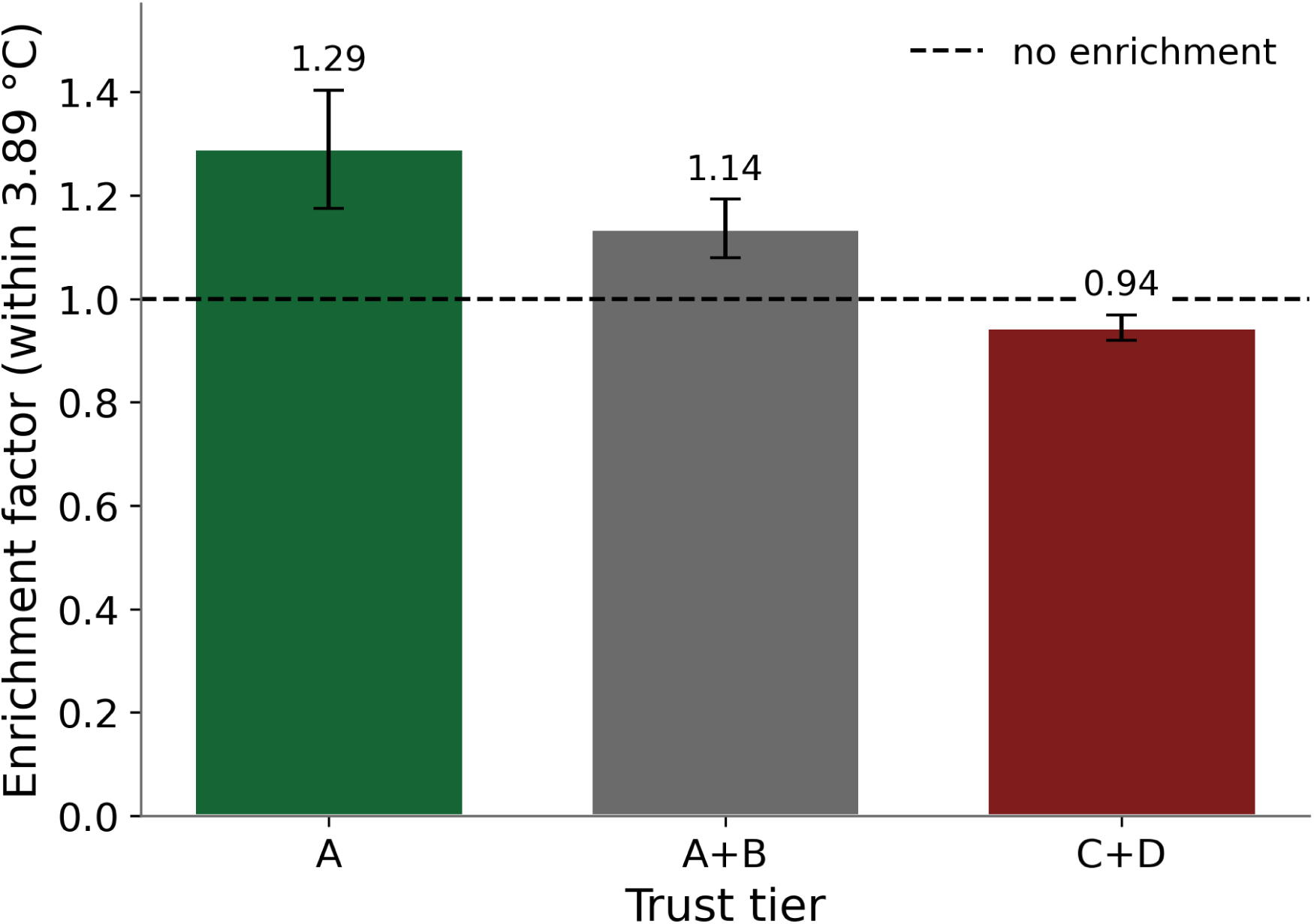
Enrichment factor for predictions within the reproducibility floor, with 95 percent bootstrap intervals; the trusted and flagged tiers exclude one in opposite directions.

The most useful comparison for this error is not against another model’s accuracy but against the measurement’s own reproducibility floor. Scenario A’s 2.99°C and the combined trusted tier’s 3.47°C both sit below the 3.89°C reproducibility floor measured from replicate assays of identical sequences, so trusted predictions are accurate to within the assay’s own noise. Expressed as an enrichment factor, the standard currency for a screening filter, Scenario A reaches 1.29 with a confidence interval of 1.18 to 1.40 while the flagged tier reaches 0.94 with an interval of 0.92 to 0.97, so both intervals exclude one in the expected directions.

However, the within-scenario Spearman correlation must be read with care. It is inverted, at 0.391 for Scenario A against 0.665 for Scenario D, which reads as though trusted predictions correlate worse. This is range restriction and not a broken axis, because Scenario A’s measured temperatures have a standard deviation of 6.24°C against Scenario D’s 10.76, and a rank correlation computed inside a compressed label range is not comparable to one computed across a wide range. The measured-Tm standard deviation column in Table 2 is printed alongside for exactly this reason. Error metrics are the primary evidence, and the per-scenario spread is reported alongside any correlation.

### 3.2 Explainability as a trust tool

Attributions are used to explain each prediction, and to be certain that attribution mass on the buried core is a good measure of explanation quality, we tested it rather than assumed it. The alternative from our own lineage, the prediction-explanation direction criterion, does not transfer to this property. As a per-protein consistency rate it reaches 0.4506 across the full dataset, below the 0.50 chance level with a p-value of 9.8 × 10^-45^, and it is below chance in every partition and every stratum tested (Table S7, Figure S1). Substituting it for the buried-core axis while holding everything else fixed gives an error separation of 0.962 times, a slight inversion, against the buried-core axis’s 1.119 times, and it reverses the ordering of Scenarios A and B. This has a measurable mechanism, since melting temperature is a global emergent property of an entire fold, so occluding a single residue barely moves the prediction and the sign of that change is close to noise, and consistent with this, 61.9 % of proteins carry a positive mean signed attribution regardless of their temperature, a systematic occlusion bias that mechanically caps the consistency rate below 0.50 for the below-mean half of the data. The direction criterion works in the transition-metal setting because properties such as orbital energies and metal partial charge are dominated by a few atoms (Onawole, 2026a), which is not the case here, and stratifying by distance from the dataset mean rules out the obvious alternative explanation, since consistency is worst furthest from the mean at 0.395 rather than nearest it, so the criterion fails everywhere rather than at a decision boundary. The buried-core axis is grounded in an established structural correlate of protein thermostability (Paiardini et al., 2008; Glyakina et al., 2007), and it clears the empirical bar as well. The established criterion from the same lineage carries no signal for this property, while the protein-specific alternative separates error in the correct direction.

Occlusion costs one forward pass per residue per model, so running it on the full ensemble was not affordable and one checkpoint was used, selected by cross-validation score alone with no reference to held-out performance. The entire trust pass was repeated once per deployed member over the identical calibration cohort and the identical held-out test set (Table S7), and the ordering of error across scenarios held in all six arms, with the trusted against untrusted separation at 1.255 times throughout and not one protein moving between the trusted and untrusted bands. That last invariance is structural rather than lucky, because the consensus axis reads the same five models whatever performs the occlusion and therefore fixes both band totals, so every disagreement between arms is a reallocation within a band and never a change of decision. What does move is the finer diagnosis, since an individual protein’s assignment to Scenario A rather than B is not stable across checkpoints, which is why per-residue attributions are reported as population-level evidence about classes of proteins rather than as a claim about any single one.

The taxonomy is most useful read as a diagnostic instrument, which is how it was used in our transition-metal work, where a per-metal breakdown of scenario rates identified a missing input feature (Onawole, 2026a). Because the taxonomy is two crossed binary axes, the informative comparisons hold one axis and vary the other, and a property counts as a driver of an axis only when it replicates in both of that axis’s matched pairs. Run across the full 20,183-protein pass, this gives each axis a mechanism (Figure 4, Table S4). Before reading design trends off the full dataset, one check matters. The scenario mix of the training pool and of the held-out test set are nearly identical, at 8.98 against 8.28 percent Scenario A (Table S1), so the trust axes are not detecting training memorisation, and the trends below are not an artefact of training exposure. Contact density drives coherence and only coherence, replicating on both coherence pairs at *p* below 10^-244^ and absent on one consensus pair at *p* equal to 0.90. Sparsely packed proteins receive high coherence. Length drives consensus, with longer proteins producing tighter ensemble agreement. Higher measured temperature lowers consensus, which is the finding that matters most for what follows, and it is confirmed independently by the organism level, since *Picrophilus torridus* is 40.7% Scenario C at 1.90 times enrichment with 0.3% in each of A and B. Thermophiles concentrate in the underconfident tier.

**Figure 4.**
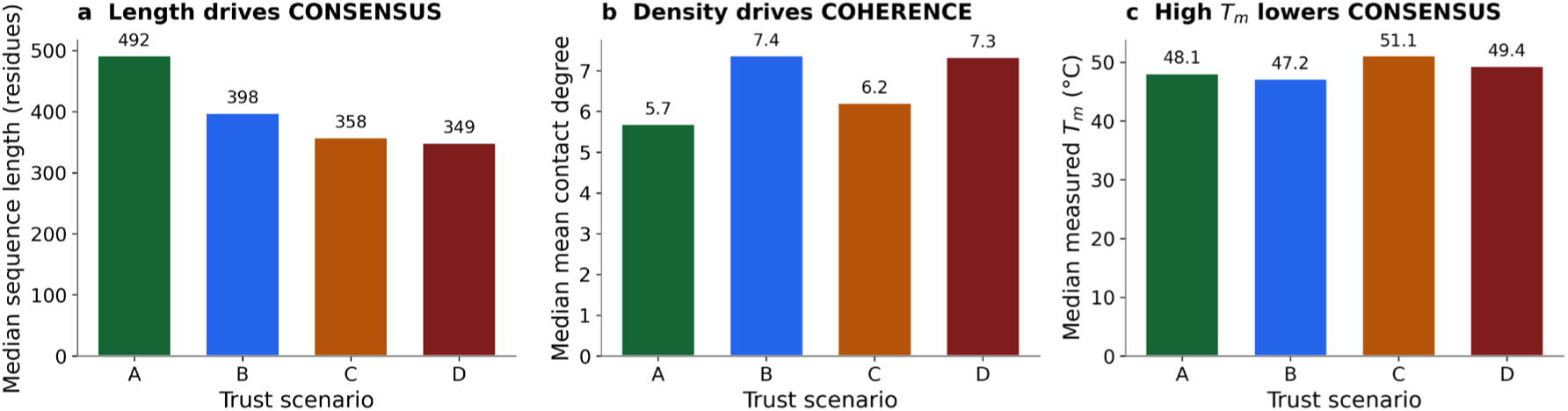
Drivers of trust axis over the full 20,183-protein pass. Medians by scenario for (a) sequence length, (b) mean contact degree and (c) measured melting temperature. Contact density drives coherence and only coherence; length drives consensus; higher melting temperature lowers consensus, which is why thermophiles concentrate in Scenario C. Design trends, not reliability estimates.

Two honest qualifications belong with this, at 20,183 proteins statistical significance is cheap, and effect sizes must lead, since contact degree on one consensus pair moves from p equal to 0.012 in the test set to 2.7 × 10^-26^ at full power on a median difference of only 5.70 against 6.21. And part of the Scenario A signature is coupling between the metrics and protein geometry rather than model reasoning, since the coherence axis correlates with mean contact degree at −0.306. The taxonomy therefore describes protein size and packing as well as model confidence, which bounds any claim that the model understands Scenario A proteins and makes stratified calibration the highest-value improvement available at no training cost. The consequence for the industrial question follows directly. The ensemble is least confident precisely on the thermostable enzymes the industrial question is about, a diagnosis with an obvious remedy, namely targeted acquisition of thermophile data, and it is Scenario C that identifies it.

The graph carries backbone contacts and a frozen sequence embedding, and it carries no cofactor, no bound metal, no oligomeric state and no solvent context. Enzymes concentrating in the low-coherence tiers and oxidoreductases reaching 65.9% Scenario D is the signature a missing cofactor representation would produce, so this was tested at full power, where 109 Pfam domains clear 25 proteins against exactly one in the test set alone. Raw rates are confounded, because being an enzyme, being short and being densely packed are each independently enriched in the refused tier, so the primary analysis is a logistic regression of each axis on a cofactor flag with enzyme status, log length and mean contact degree in the model. The pre-specified pooled flag gives an odds ratio for Scenario D of 1.29 with an interval of 1.15 to 1.44 (Table S1). That pooled figure understates the effect, because it averages two opposite classes.

Split apart, the contrast is the strongest mechanistic result here (Figure 5). Redox cofactors, covering heme, flavin, NAD(P), iron-sulfur clusters, quinone and molybdenum, give an odds ratio of 1.66 with an interval of 1.42 to 1.93, while structural metal sites are null at 0.91 with an interval of 0.75 to 1.10. Redox-cofactor proteins reach 2.9% Scenario A and 66.9% Scenario D against base rates of 8.9 and 48.9%, whereas Cys2-His2 zinc fingers reach 26.9% Scenario A, three times the base rate, and are the most trusted class in the dataset.

**Figure 5.**
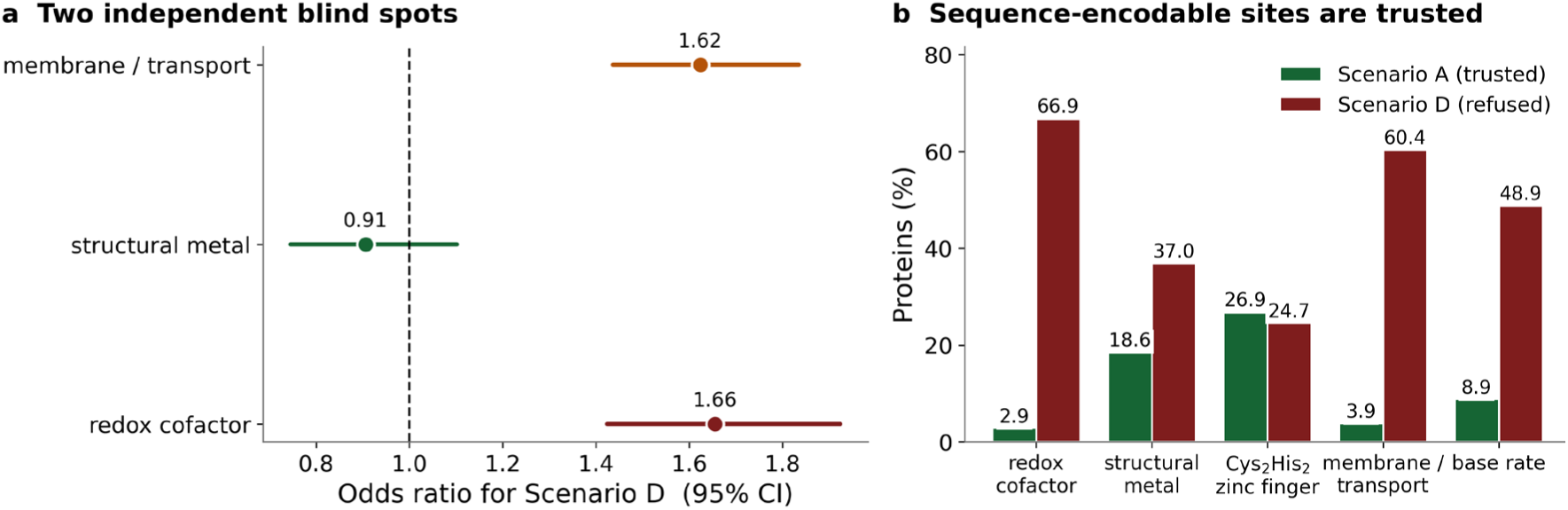
Two independent representational blind spots, over the full 20,183-protein pass. (a) Odds ratios for the refused tier from a logistic model controlling for enzyme status, log length and mean contact degree. (b) Trusted and refused rates by class. Redox cofactors depress reliability while structural metal sites do not, and Cys2His2 zinc fingers, whose metal site is a recoverable sequence motif, are the most trusted class in the dataset.

The mechanism is what makes this a representational claim rather than a correlation. A Cys2His2 zinc finger’s metal site is a sequence motif with canonical spacing, so the frozen embedding encodes it directly and the model is at its most reliable there. A heme, flavin or iron-sulfur cluster is held by residues dispersed across the fold, and neither its identity nor its occupancy is recoverable from sequence or from a backbone contact map. The same nominally missing input is harmless when sequence can recover it and damaging when it cannot, which is a positive control sitting inside the hypothesis. Every redox class moves the same way, and the effect survives restriction to enzymes alone.

A second blind spot was found by reading the ranked domain list rather than by prediction. Three transporter families sit near the top of the refused tier while being no percent enzyme and only 3 to 7% cofactor-flagged, so they cannot be the cofactor effect relabelled. Membrane and transport proteins reach 3.9% Scenario A and 60.4% Scenario D, and in a joint model both effects survive, with odds ratios of 1.62 for membrane and 1.67 for redox on Scenario D, overlapping in only 29 proteins. These are two independent failure modes of comparable size. Their mechanisms differ, and the difference bounds what any architectural fix can buy. An AlphaFold model carries no membrane, so the contact graph describes an integral membrane protein as though it were soluble and its hydrophobic surface reads as though it were misfolded. Separately, a lysate-based melting curve for a membrane protein is not measuring the same physical event it measures for a soluble protein. This blind spot is therefore part representation and part label validity, and only the first part is addressable by adding features. Two limitations travel with these results. Cofactor status is inferred by keyword matching on protein names rather than from a curated annotation, so recall is low and every effect above is a lower bound. The split between redox and structural classes is post-hoc, and the pooled odds ratio of 1.29 is the pre-specified figure.

### 3.3 External dataset evaluation

The PET hydrolase set was used to test whether the model is practically useful on proteins it never trained on, assayed by differential scanning fluorimetry (DSF) on purified protein rather than by the mass-spectrometry-based method used for training. On the 233 PET enzymes with a measured melting temperature, the model reached a Spearman correlation of 0.2802 with an interval of 0.144 to 0.403, a root mean squared error of 10.44°C, and only 0.37°C of error saved relative to always predicting the training mean, an interval that includes zero (Table 3). Predictions are compressed by a factor of 2.49 relative to the measured spread and carry a bias of −4.69°C. Any statement about accuracy on this set must say whether it concerns the ranking or the scale, because the two give opposite answers, the model ranks PET enzymes better than chance while retaining almost none of its internal skill over a constant predictor.

**Table 3.**
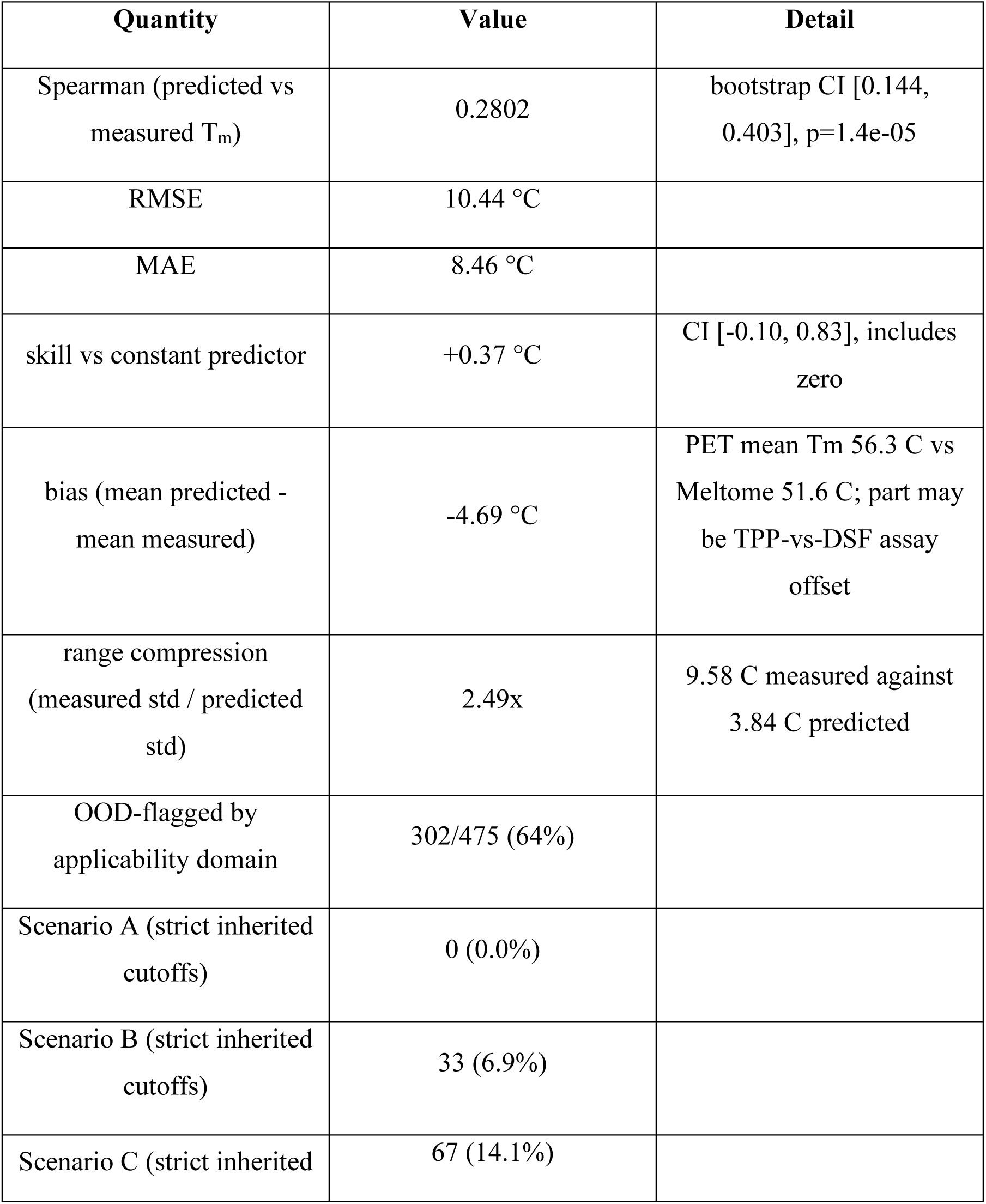
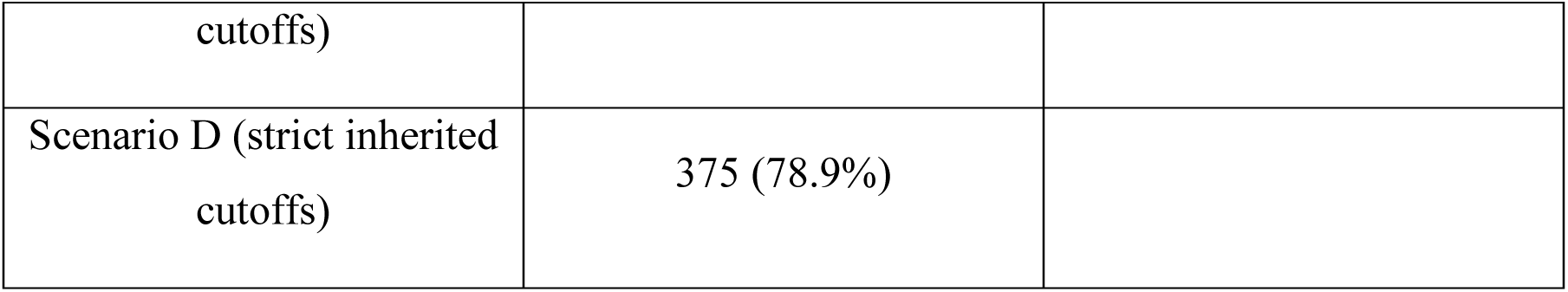
Transfer to the 475-enzyme PET hydrolase set. 233 enzymes carry a measured melting temperature.

Under the inherited cutoffs Scenario A is empty and 78.9% of enzymes fall into the refused tier (Figure 6), a degenerate labelling that is substantially a calibration-transfer artefact rather than a competence verdict. The distribution-shape diagnostic places this set in the rescuable category, since removing location and scale collapses the discrepancy between distributions almost entirely, with a Kolmogorov-Smirnov statistic on the consensus axis falling from 0.453 to 0.041 and a quantile-quantile coefficient of determination of 0.987. Applying the same percentile ranks within the external cohort restores a non-degenerate labelling at 3.0% Scenario A, against a pre-registered blind prediction of around 4%. One exclusion is worth stating plainly, since it flatters these numbers. The benchmark hydrolase LCC-ICCG and the expression control Red Fluorescent Protein (RFP) are excluded throughout as non-candidates. LCC-ICCG carried the highest measured Tm in the spreadsheet at 93.6°C and was the single largest prediction error, so removing it improves every accuracy metric here. Over all 475 rows the same quantities are Spearman 0.2748, RMSE 10.79°C, MAE 8.61°C, skill +0.37°C and range compression 2.57x. The exclusion rule is about provenance, not residual size.

**Figure 6.**
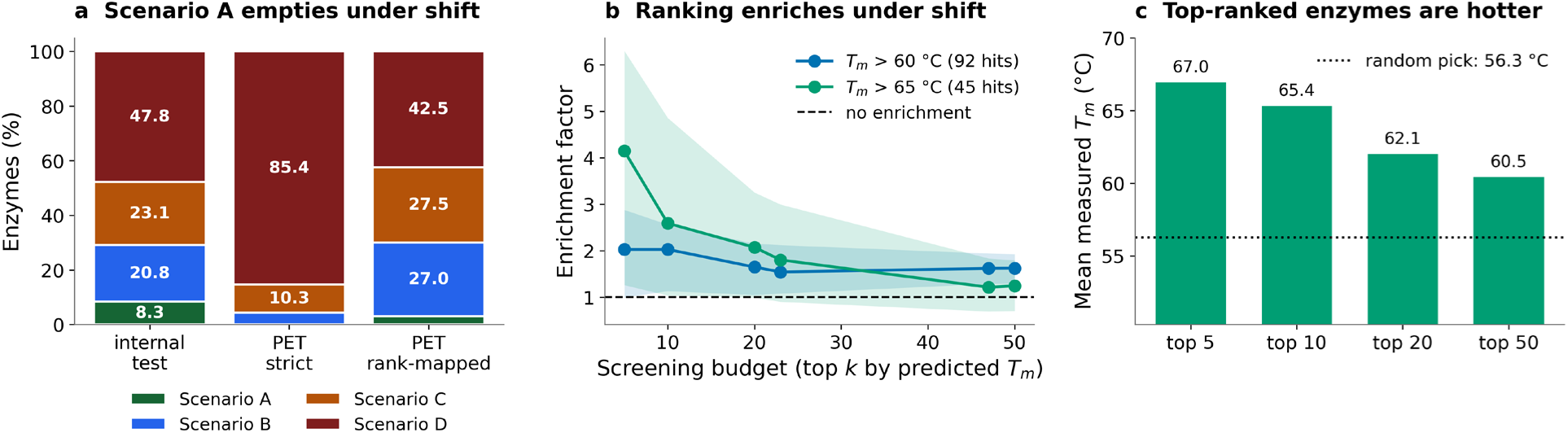
Transfer to 475 PET hydrolases at a median 10 % sequence identity. (a) Scenario mix for the held-out test set and for the external set under strict inherited cutoffs and under rank-mapped cutoffs. (b) Enrichment factor against screening budget at the two thresholds the labelled set can support, with bootstrap intervals. (c) Mean measured melting temperature of the top-ranked selection against a random pick.

One confound should be stated before the bias is read as a model failure. The external set’s mean measured temperature exceeds the training mean by 4.6°C, and the observed bias is −4.69°C. Training labels come from mass-spectrometry-based melting curves on lysate, and external labels from differential scanning fluorimetry on purified protein, so a systematic assay offset would produce this signature. This does not affect the rank correlation.

A control refutes the obvious alternative explanation. Scenario A internally is long and sparsely packed while these enzymes are short and compact, so the collapse could be the metric coupling described earlier re-reading protein size. Training proteins matched to the external length range still earn Scenario A about 7% of the time, against zero for the external set, and both axes still shift strongly after controlling for length. The collapse reflects genuine family novelty on top of any size effect.

What survives is the ordering, and it is the practically useful result. Ranking the 233 labelled enzymes by predicted temperature, the 50 highest-ranked recovered 32 of the 92 enzymes above 60°C, an enrichment of 1.62 with an interval of 1.29 to 1.92 and a p-value of 7.1×10^-5^, while the ten highest-ranked reach an enrichment of 2.59 above 65°C (Table 4, Figure 6). Enrichment decays monotonically with the size of the selection, the expected shape for a genuinely front-loaded ranking. The mean measured temperature of the top five is 67.0°C against a set mean of 56.3°C.

**Table 4.**
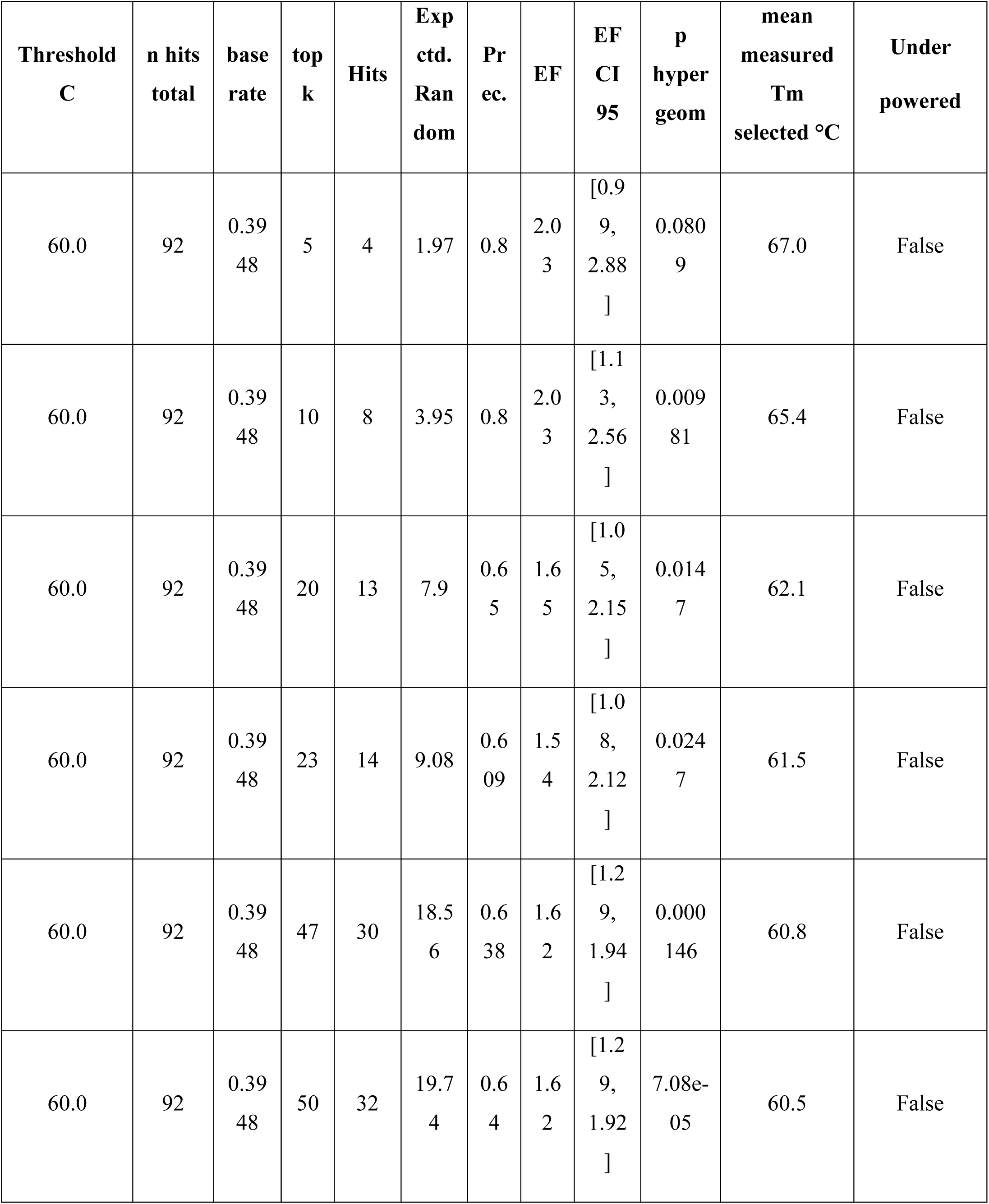
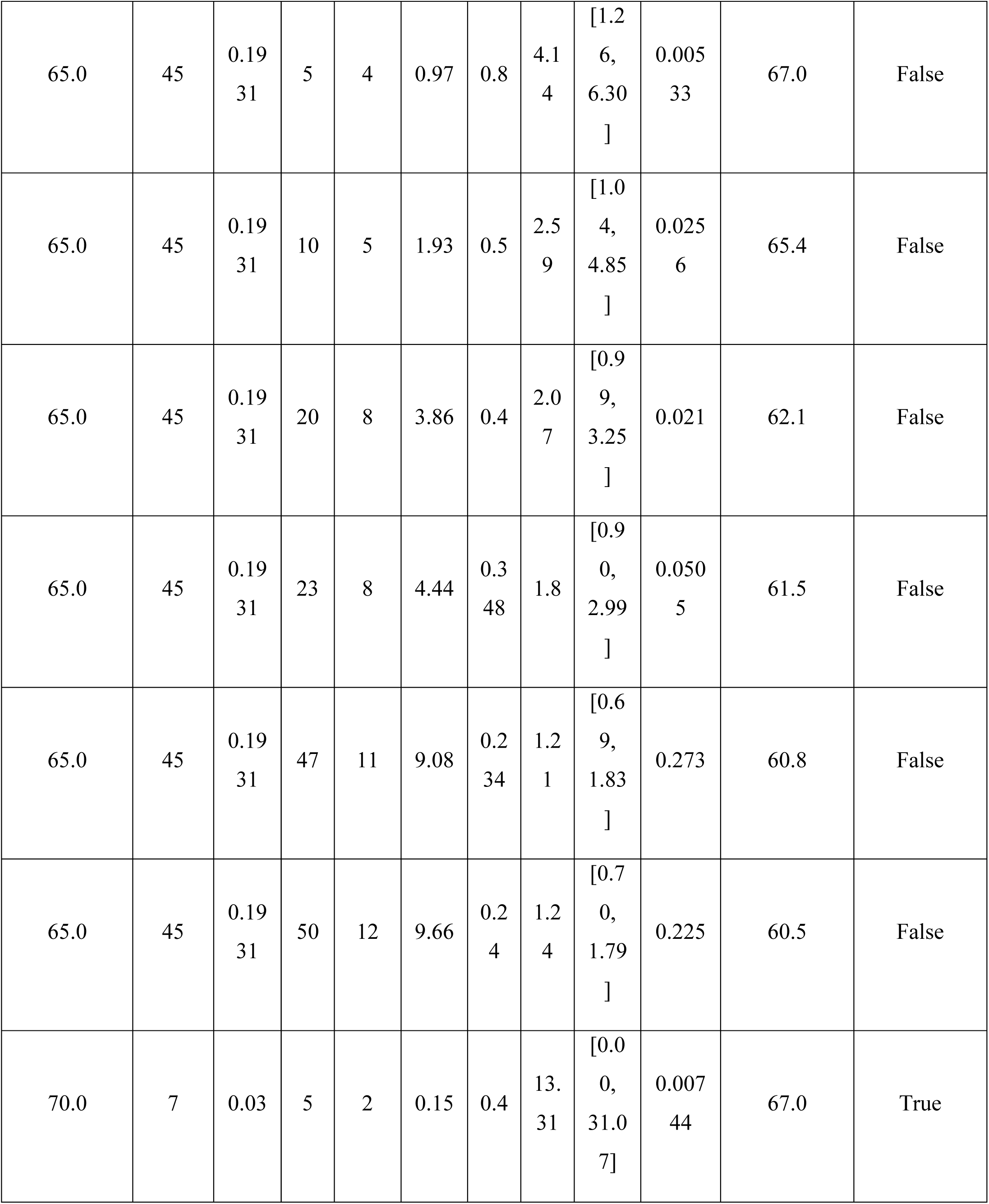
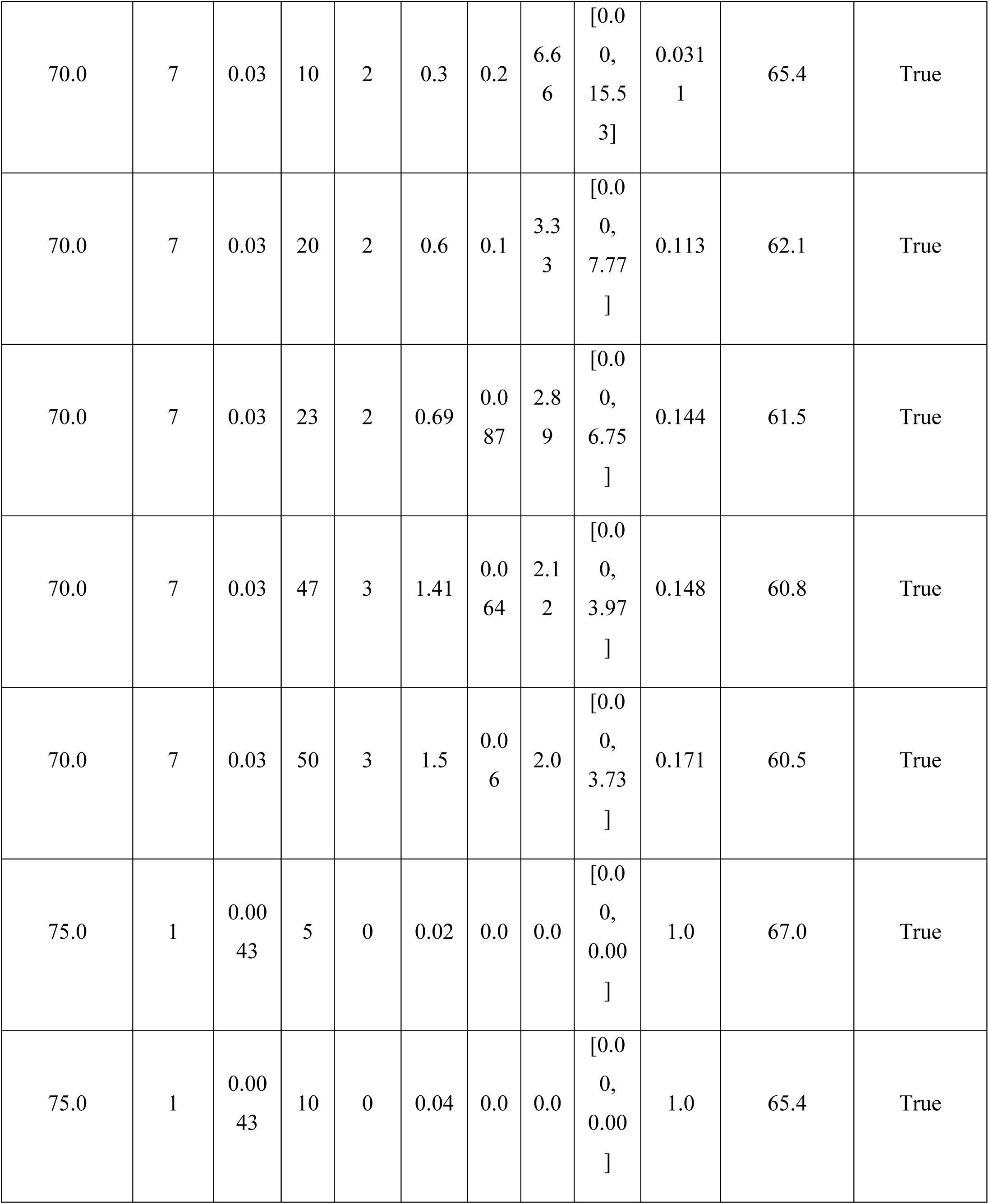
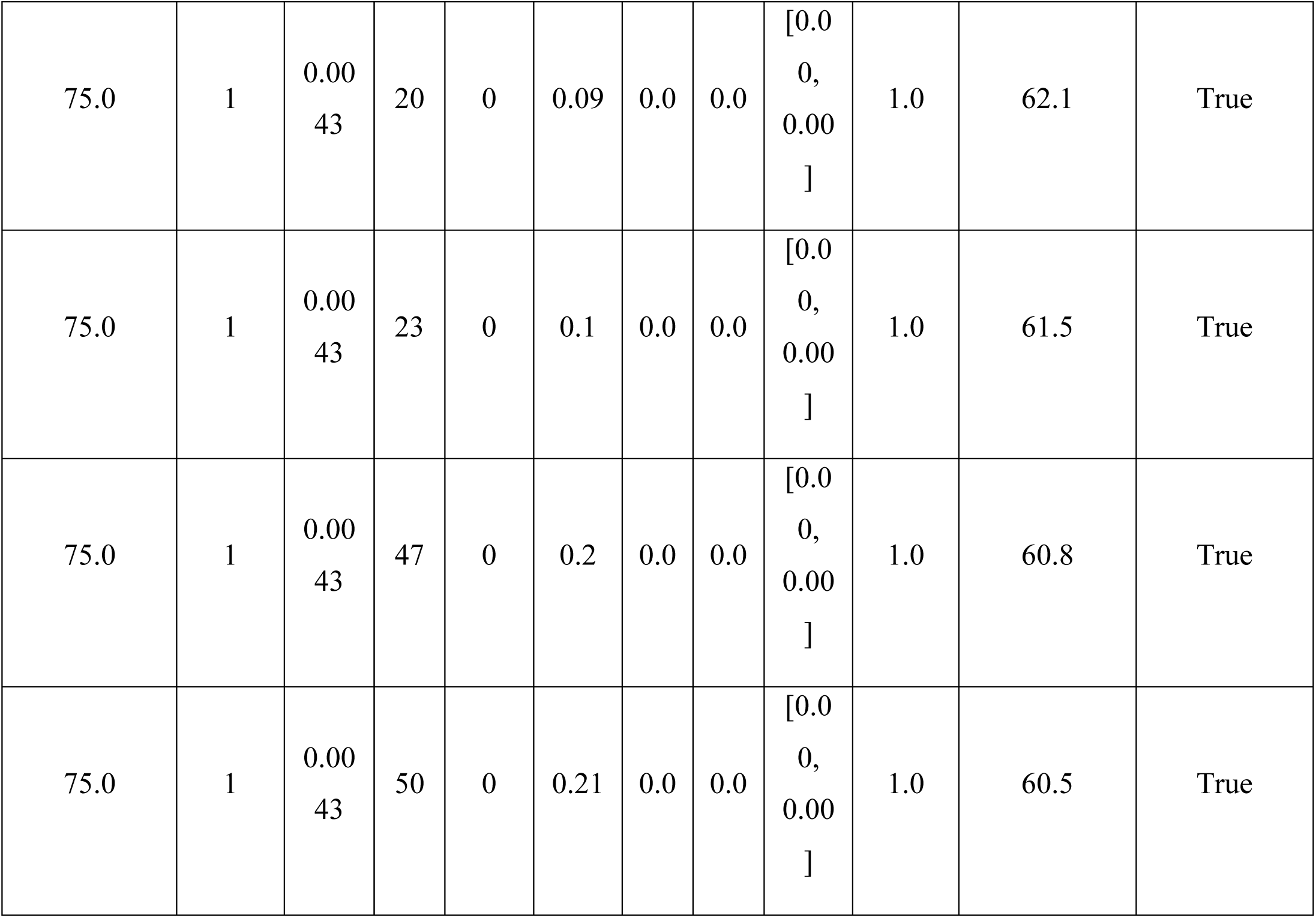
Screening enrichment on the 233 Tm-labelled PET enzymes. Enrichment factor (EF) is precision at *k* divided by the base rate; *p* is an exact hypergeometric test against a random pick of *k*; intervals are 2,000-fold bootstraps in which each replicate re-ranks and uses its own base rate.

This is the deliverable the external evaluation rests on. Instead of expressing and assaying all 233 enzymes, testing the top 50 recovers 32 of the 92 enzymes above 60°C, an enrichment of 1.62 with an interval excluding 1 (p=7.1×10^-5^). Three things travel with this table. The 70°C and 75°C thresholds are underpowered, 7 and 1 positives respectively, and are printed only so the choice of 65°C as the highest claim-bearing threshold is documented; Tm > 75°C selects zero hits because the predicted maximum is 58.7°C against a measured maximum of 82.3°C, which bounds the claim to moderately rather than extremely thermostable enzymes. The top-5 selection is the same five enzymes at every threshold, so those rows are not independent findings. Where the interval and the p-value disagree, the interval is the one to trust, since it carries base-rate noise the hypergeometric test conditions away.

This claim has a ceiling, above 75°C the ranking recovers nothing, because the model’s maximum prediction is 58.7°C against a measured maximum of 82.3°C, so range compression reappears in the screening currency. The ranking finds moderately thermostable enzymes, not extreme ones, matching the framing that stability determines evolutionary potential and that the practical task is ranking starting points by stability headroom rather than predicting an absolute value [26].

Three further experiments identify the cause of this collapse. First, the applicability-domain axis inverts. The model is measurably better on enzymes that axis calls out of domain, with a skill difference of −1.83°C and an interval of −2.93 to −0.72. The mechanism is that far-from-training enzymes receive mean-reverted predictions, and mean reversion is a safe strategy when the label spread is wide, so the axis identifies unfamiliarity correctly and then labels the safe, accurate predictions as untrustworthy. The same inversion appears independently in the industrial decision, where adding the in-domain filter reduces the candidate pool from 53 to 12 and its hits from three to zero (Table S8). The axis must be rank-based rather than value-based before it is ever used to steer generation.

Second, whether the structure encoder earns its place over a simpler baseline can now be tested on both cohorts together. A ridge regression on mean-pooled ESM2 embeddings was trained on the same pool with the same cluster splits and scored on both the internal test set and the external PET set, paired on identical proteins (Table 5). Internally the RGCN is better on both error and ranking (Spearman 0.6476 versus 0.5735, skill 3.03°C versus 2.41°C), so the structure encoder earns its place there. Externally the result splits, the ridge achieves better absolute-scale skill because it is less biased (skill 1.08°C versus 0.37°C), while the RGCN still ranks better (Spearman 0.2802 versus 0.2356). All four paired intervals exclude zero, a genuine double dissociation. Both models lose most of their skill under family-level shift, the RGCN retaining 12% of its internal figure and the ridge 45%. In summary, both architectures collapse, but they collapse differently, when evaluated on an out of domain external dataset.

**Table 5.**
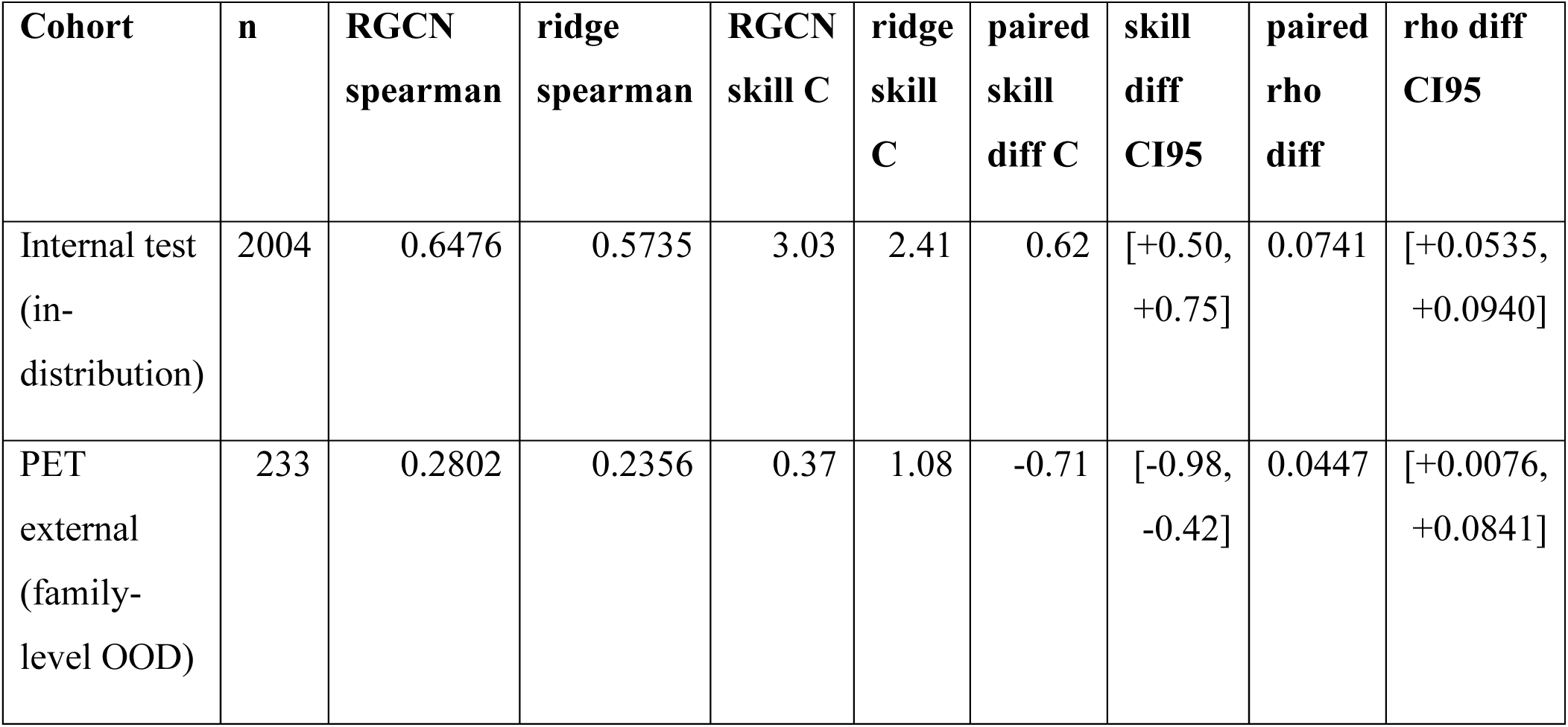
Paired accuracy against the ridge baseline. Skill is error saved relative to the training-pool mean (51.63°C). Intervals are 2,000-fold bootstraps over proteins, resampling both models on the same index set.

Third, one experiment isolates taxonomy as the cause directly. All prokaryotes were removed from training, and the model was retrained from scratch, with the assay, the structure source and the pipeline held constant. Scenario A fell from 11.4% in the size-matched random control to 0.9% (Figure 7, Table S5). The external collapse therefore needs no external-set-specific explanation, since it requires neither 10% sequence identity, nor the assay change, nor predicted rather than deposited structures. Any distribution shift does this.

**Figure 7.**
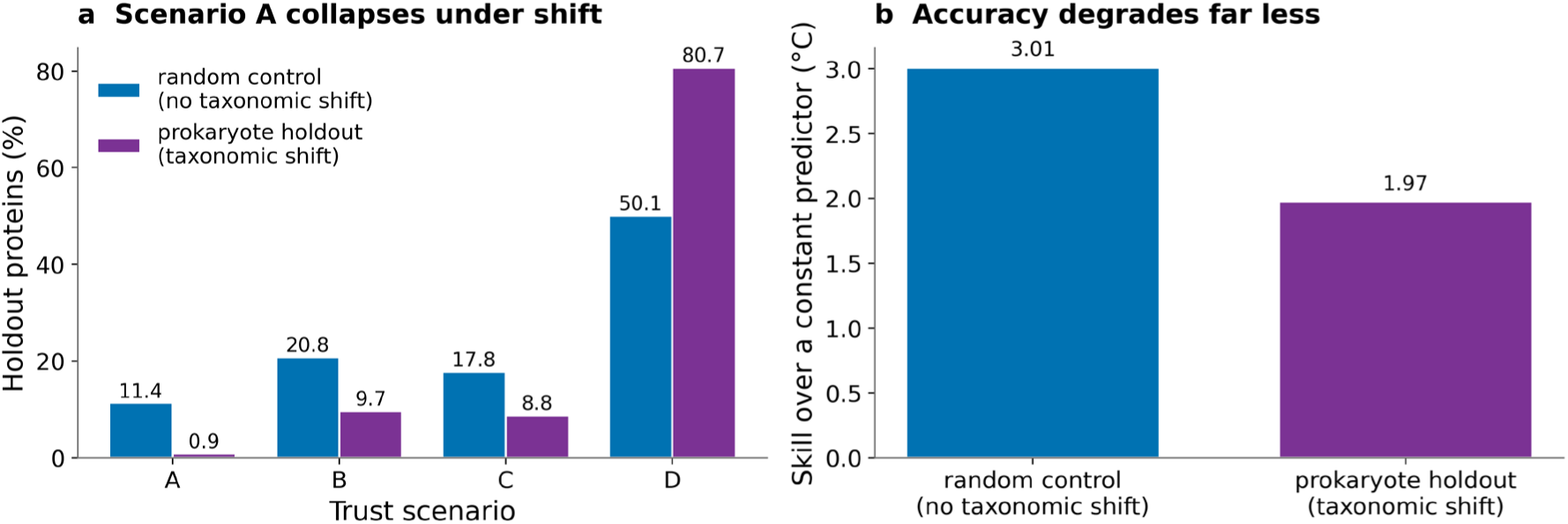
Leave-one-taxon-out ablation. All prokaryotes were removed from training, with the assay, the structure source and the pipeline held constant, against a size-matched random holdout control. (a) Scenario coverage; the trusted tier collapses under a taxonomy-only shift. (b) Error saved relative to a constant predictor. Both arms are retrained from scratch, and neither is the deployed ensemble.

The marginal enrichment of 1.30 (interval 1.14 to 1.47) for the trusted tier is largely a composition effect rather than per-protein discrimination. The five held-out organisms differ enormously in difficulty, and testing within organism instead gives an enrichment of only 1.12, with an interval spanning one, carried almost entirely by a single species. The framework still works decisively at the level of a population, refusing to vouch for 0 of 177 *Picrophilus* and 6 of 521 *Thermus* proteins, but its per-protein effect is real and small rather than large, around 1.05 in distribution. A unifying mechanism is visible in both settings. The model cannot extrapolate above its training temperature range, and *Thermus* at a median 85.6°C is simply unreachable, giving a mean absolute error of 26.7°C. The external set shows the same failure as range compression and negative bias. One residual confound should be noted, the prokaryote arm trained on about 5% fewer records than its control because of leakage-driven cluster removal.

### 3.4 What this means for deployment

The results support a conditional rule rather than a single recommendation, and the condition is how far the query sits from training. Inside the competence domain, prioritise by trust label, which enriches at 1.29 and does so on both model classes tested. Outside it, prioritise by rank, because the trust label there is a refusal signal rather than a triage tool while the ranking still enriches at 1.62. A practitioner can act on that. Two qualifications keep the rule honest. Consensus is the stronger axis and is what should drive screening, with coherence retained for mechanistic interpretation at the level of a class of proteins, namely the two blind spots, rather than for triage or for a claim about any single protein. And whether the trusted-tier label itself carries meaning this far from training remains open, since the external set was deliberately chosen to be out-of-distribution.

## 4. Conclusion

We present ProtTrust-XAI, a three-axis reliability framework for protein property models, and measure where the competence of a structure-grounded thermostability model ends. The competence boundary, rather than an accuracy figure, is what this work delivers, because a model that reports where it stops being trustworthy can be used by someone who did not build it. Two results give that boundary substance. Used as a diagnostic instrument rather than a reporting category, the taxonomy localised two independent representational blind spots and supplied a mechanism for each, with redox cofactors depressing reliability while structural metal sites do not and sequence-encodable zinc fingers emerging as the most trusted class in the data. Under family-level shift the absolute temperature scale collapsed while the ranking survived, which a controlled taxonomy-only ablation reproduced on the training data, so the failure is a property of distribution shift rather than of the external set. The framework refuses to vouch for whole taxa it cannot handle, which is what it exists to do.

Four limitations bound these claims, and each points to a specific next step. The coherence axis rests on an established structural correlate of stability rather than a validated causal explanation [23, 24], so the highest-value fix is calibrating trust percentiles within size and density strata, which needs enough data per stratum to calibrate reliably and a re-validation of everything built on the current calibration, which is why it is future work rather than a correction made here. The graph carries no cofactor, ligand, disulfide or oligomeric information, the leading explanation for the two blind spots, so adding those features is a targeted repair concentrated in the enzyme classes already diagnosed. The applicability-domain axis inverted under shift and must be made rank-based before it constrains generation, since as implemented it would steer a design loop away from its most accurate predictions. The external set, though deliberately chosen to be out-of-distribution, is too small, 21 of 233 labelled enzymes reaching the trusted tier, to resolve whether that label transfers, so a future benchmark needs the trusted tier alone to reach a hundred labelled examples. The plastic-degradation case study adds three further limits. Firstly, the predictions compress toward the mean, so the ranking recovers moderately thermostable hydrolases rather than the most extreme ones. Secondly, the candidate structures were predicted with a different method from the training structures, a stated caveat rather than a measured effect. Finally, the fully characterised subset is too small to claim significance on the expressed-stable-active endpoint that matters industrially, though the condition-resolved activity data needed to sharpen that decision already exists, and mutation-level stability data would extend the framework from choosing a starting point to choosing a substitution.

Because the three axes read ensemble behaviour, attribution geometry and distance from the training data, none of which reference melting temperature, the construction transfers to any protein property for which an ensemble and a per-residue attribution can be computed, and the settings it transfers to are ones where acting on a wrong prediction is expensive. Enzyme technology for food and beverage production is the nearest case, where stability sets the temperature a process can run at and where a fixed budget must decide which candidates to express and assay. Biologics development poses the same problem for different endpoints, since solubility, aggregation propensity and expression yield are predicted for antibody candidates engineered away from any natural repertoire, which is precisely the shifted regime in which a point estimate is least informative and an explicit refusal is most valuable. Immunogen and vaccine design works on sequences that are novel by intention, so a model asked about a designed antigen is being asked to extrapolate, and establishing whether it can is a precondition for using its answer. Plastic depolymerisation, the case study here, sits in the same family of problems and was chosen because it was able to stress the framework. In each of these settings, the deliverable is not a better average accuracy but a defensible decision about which predictions to act on, and measuring where a model stops being trustworthy is the prerequisite for designing inside that boundary.

## Acknowledgement

This work was supported by resources provided by The University of Queensland Research Computing Centre’s Bunya supercomputer.

## Authors contribution

**Abdulmujeeb O. Onawole**: Conceptualization, Methodology, Writing - review and editing. **Rukayat O. Adegoke**: Conceptualization, Writing - review and editing. All authors read and approved the final manuscript.

## Data and code availability

All code needed to reproduce the pipeline, from data preparation and graph construction through cross-validation, occlusion attribution, trust calibration and the external evaluation, is available at https://github.com/MujeebOnawole/prottrust-xai. An interactive webtool is available at https://huggingface.co/spaces/catenate/prottrust-xai.

