## Supplementary material for "Mapping the Competence Boundary of a Protein Property Model: A Case Study on Plastic-Degrading Enzymes using ProtTrust-XAI": Supplemntal Information

This document collects the supporting tables and figures referenced in the main text. Two conventions apply throughout and are stated here. First, every reliability figure, meaning any scenario coverage, error separation or enrichment estimate quoted as evidence of trustworthiness, is computed on the held-out test set. Second, tables reporting design trends are computed over the full 20,183-protein pass and say so in their captions; because most of those proteins are training data, no reliability claim may be drawn from them.

### S1. Partition and calibration

**Table S1. Trust-scenario coverage broken down by data partition, over the full 20,183-protein pass. Requested because a scenario distribution computed over training data would overstate real-world reliability.**

| **Partition** | **n** | **A** | **B** | **C** | **D** |
| --- | --- | --- | --- | --- | --- |
| pool | 18179 | 8.98 | 20.77 | 21.24 | 49.0 |
| test | 2004 | 8.28 | 20.81 | 23.1 | 47.8 |

The pool and held-out test mixes are nearly identical (Scenario A 8.98% against 8.28%), so the trust axes are NOT detecting training memorisation. Every reliability number in the main text is nonetheless taken from the test partition alone.

**Table S2. Architecture and optimiser settings selected by 40 Optuna trials on a family-level cluster fold, pre-registered before cross-validation. The contact-distance cutoff was fixed a priori at 8.0 Angstrom because searching it would require rebuilding roughly 20,000 graphs per value.**

| **parameter** | **value** |
| --- | --- |
| lr | 0.00035119753147402075 |
| weight_decay | 0.005669287259919309 |
| batch_size | 16 |
| num_layers | 2 |
| node_dim | 128 |
| dropout | 0.05092185149909252 |
| CV Spearman (5 folds x 3 repeats) | 0.6554 +/- 0.0126 |

The depth search returned a clean negative result rather than a null one: two layers won, and every six-layer trial sits at the bottom of the table, which is a measured over-smoothing signal. The whole landscape spans only about 0.02 Spearman.

Early stopping, not dropout, is the effective capacity control: the best epoch is between 4 and 8 in every one of the 15 runs.

### S2. Design-level analyses over the full dataset

*These analyses require the whole dataset rather than the held-out split, because the per-category counts that the diagnoses depend on are not reachable at test-set size. Exactly one Pfam domain clears 25 proteins in the test set against 109 at full power. Design trends only; not reliability estimates.*

**Table S3. Logistic regression of trust scenario on cofactor dependence, over the full 20,183-protein pass, controlling for enzyme status, log protein length and mean contact degree. Design analysis, not a reliability claim.**

| **Specification** | **Outcome** | **Term** | **Odds ratio** | **C.I. 95** | **p** |
| --- | --- | --- | --- | --- | --- |
| Pooled | low_coherence (B or D) | cofactor | 1.191 | [1.04, 1.36] | 0.00998 |
| Pooled | low_coherence (B or D) | contact_degree | 1.499 | [1.47, 1.53] | 3.3100000000000004e-271 |
| Pooled | low_coherence (B or D) | is_enzyme | 1.803 | [1.66, 1.95] | 7.38e-47 |
| Pooled | low_coherence (B or D) | log_length | 0.517 | [0.46, 0.58] | 7.020000000000001e-27 |
| Pooled | low_consensus (C or D) | cofactor | 1.17 | [1.03, 1.32] | 0.0127 |
| Pooled | low_consensus (C or D) | contact_degree | 1.069 | [1.05, 1.09] | 1.23e-09 |
| Pooled | low_consensus (C or D) | is_enzyme | 0.915 | [0.85, 0.98] | 0.015 |
| Pooled | low_consensus (C or D) | log_length | 0.307 | [0.27, 0.35] | 1.41e-78 |
| Pooled | scenario D (both) | cofactor | 1.286 | [1.15, 1.44] | 1.27e-05 |
| Pooled | scenario D (both) | contact_degree | 1.309 | [1.28, 1.34] | 1.81e-134 |
| Pooled | scenario D (both) | is_enzyme | 1.245 | [1.16, 1.33] | 1.01e-10 |
| Pooled | scenario D (both) | log_length | 0.381 | [0.34, 0.43] | 1.21e-64 |
| Split | low_coherence (B or D) | redox_cofactor | 1.747 | [1.42, 2.14] | 8.78e-08 |
| Split | low_coherence (B or D) | structural_metal | 0.811 | [0.67, 0.99] | 0.0383 |
| Split | low_coherence (B or D) | contact_degree | 1.491 | [1.46, 1.53] | 1.61e-261 |
| Split | low_coherence (B or D) | is_enzyme | 1.783 | [1.64, 1.93] | 4.3599999999999996e-45 |
| Split | low_coherence (B or D) | log_length | 0.536 | [0.48, 0.61] | 7.989999999999999e-24 |
| Split | low_consensus (C or D) | redox_cofactor | 1.392 | [1.17, 1.65] | 0.000156 |
| split | low_consensus (C or D) | structural_metal | 0.948 | [0.78, 1.15] | 0.593 |
| split | low_consensus (C or D) | contact_degree | 1.066 | [1.04, 1.09] | 9.8e-09 |
| split | low_consensus (C or D) | is_enzyme | 0.91 | [0.85, 0.98] | 0.00975 |
| split | low_consensus (C or D) | log_length | 0.312 | [0.28, 0.35] | 6.03e-76 |
| Split | scenario D (both) | redox_cofactor | 1.656 | [1.42, 1.93] | 5.56e-11 |
| Split | scenario D (both) | structural_metal | 0.907 | [0.75, 1.10] | 0.327 |
| Split | scenario D (both) | contact_degree | 1.303 | [1.28, 1.33] | 1.8799999999999998e-128 |
| Split | scenario D (both) | is_enzyme | 1.235 | [1.16, 1.32] | 5.16e-10 |
| Split | scenario D (both) | log_length | 0.391 | [0.35, 0.44] | 9.749999999999999e-61 |
| Two_blind_spots | low_coherence (B or D) | redox_cofactor | 1.755 | [1.43, 2.15] | 6.93e-08 |
| Two_blind_spots | low_coherence (B or D) | contact_degree | 1.494 | [1.46, 1.53] | 9.2e-264 |
| Two_blind_spots | low_coherence (B or D) | is_enzyme | 1.785 | [1.65, 1.94] | 9.749999999999999e-45 |
| Two_blind_spots | low_coherence (B or D) | log_length | 0.532 | [0.47, 0.60] | 1.87e-24 |
| Two_blind_spots | low_coherence (B or D) | membrane | 1.039 | [0.91, 1.19] | 0.584 |
| Two_blind_spots | low_consensus (C or D) | redox_cofactor | 1.407 | [1.18, 1.67] | 9.69e-05 |
| Two_blind_spots | low_consensus (C or D) | contact_degree | 1.051 | [1.03, 1.07] | 8.47e-06 |
| Two_blind_spots | low_consensus (C or D) | is_enzyme | 0.964 | [0.90, 1.04] | 0.315 |
| Two_blind_spots | low_consensus (C or D) | log_length | 0.304 | [0.27, 0.34] | 9.74e-79 |
| Two_blind_spots | low_consensus (C or D) | membrane | 2.462 | [2.09, 2.90] | 2.32e-27 |
| Two_blind_spots | scenario D (both) | redox_cofactor | 1.669 | [1.43, 1.94] | 2.84e-11 |
| Two_blind_spots | scenario D (both) | contact_degree | 1.293 | [1.27, 1.32] | 2.06e-121 |
| Two_blind_spots | scenario D (both) | is_enzyme | 1.28 | [1.20, 1.37] | 6.13e-13 |
| Two_blind_spots | scenario D (both) | log_length | 0.385 | [0.34, 0.43] | 8.540000000000001e-63 |
| Two_blind_spots | scenario D (both) | membrane | 1.624 | [1.44, 1.84] | 9.73e-15 |

The pre-specified pooled cofactor flag UNDERSTATES the effect because it averages two opposite classes. Split apart, redox cofactors (heme, flavin, NAD(P), Fe-S, quinone, molybdenum) raise the odds of Scenario D at OR 1.66, while structural metal sites are null at 0.91. Cys2His2 zinc fingers are the positive control, reaching 26.9% Scenario A against a 8.9% base rate, because a zinc finger site is a sequence motif the frozen embedding already encodes whereas a heme or Fe-S cluster is recoverable neither from sequence nor from a backbone contact map.

LIMITATIONS: cofactor status is inferred by keyword match on the UniProt protein name rather than from a curated annotation, so every effect is a lower bound; the redox-versus-structural split is post-hoc, and the pooled OR of 1.29 is the pre-specified number.

**Table S4. Matched-pair contrasts isolating each trust axis, over the full 20,183-protein pass. The taxonomy is two crossed binary axes, so the informative comparisons hold one axis and vary the other: A-vs-B and C-vs-D isolate coherence, A-vs-C and B-vs-D isolate consensus. A property counts as a genuine driver of an axis only when it replicates in BOTH of that axis's pairs.**

| **contrast** | **isolates axis** | **median group1** | **median group2** | **p** |
| --- | --- | --- | --- | --- |
| A_vs_B_n_residues | coherence | 492.0 | 398.0 | 3.03e-24 |
| A_vs_B_mean_contact_degree | coherence | 5.7 | 7.376 | 3.73e-245 |
| A_vs_B_measured_tm | coherence | 48.13 | 47.222 | 6.19e-14 |
| C_vs_D_n_residues | coherence | 358.0 | 349.0 | 0.00274 |
| C_vs_D_mean_contact_degree | coherence | 6.211 | 7.338 | 1.3700000000000002e-248 |
| C_vs_D_measured_tm | coherence | 51.149 | 49.397 | 2.27e-53 |
| A_vs_C_n_residues | consensus | 492.0 | 358.0 | 4.14e-57 |
| A_vs_C_mean_contact_degree | consensus | 5.7 | 6.211 | 2.66e-26 |
| A_vs_C_measured_tm | consensus | 48.13 | 51.149 | 9.2e-75 |
| B_vs_D_n_residues | consensus | 398.0 | 349.0 | 2.97e-36 |
| B_vs_D_mean_contact_degree | consensus | 7.376 | 7.338 | 0.903 |
| B_vs_D_measured_tm | consensus | 47.222 | 49.397 | 1.78e-78 |

Contact density drives coherence and only coherence. Length drives consensus. Higher measured Tm lowers consensus, which is why thermophiles concentrate in Scenario C.

AT n=20,183 SIGNIFICANCE IS CHEAP AND EFFECT SIZE MUST LEAD. Contact degree on the A-versus-C pair moves from p=0.012 in the test set to p=2.7e-26 here on a median difference of only 5.70 against 6.21. Quote medians; do not let a p-value carry a claim.

### S3. Distribution shift and ablation

*The leave-one-taxon-out arms retrain from scratch and are therefore an ablation. No number in this section describes the deployed ensemble, which saw every organism and cannot be evaluated on any of them.*

**Table S5. Leave-one-taxon-out ablation. All 6,411 prokaryotes were held out of TRAINING, with the assay, the structure source and the pipeline held constant, so only taxonomy changes. The size-matched random control separates taxonomic shift from the mere loss of training data.**

| **arm** | **n train records** | **n holdout proteins** | **spearman** | **MAE C** | **RMSE C** | **skill vs constant C** | **coverage A** | **coverage B** | **coverage C** | **coverage D** |
| --- | --- | --- | --- | --- | --- | --- | --- | --- | --- | --- |
| random control (no taxonomic shift) | 17881 | 5209 | 0.6632 | 4.1 | 5.45 | 3.01 | 11.4 | 20.8 | 17.8 | 50.1 |
| prokaryote holdout (taxonomic shift) | 16943 | 4866 | 0.4893 | 10.13 | 13.8 | 1.97 | 0.9 | 9.7 | 8.8 | 80.7 |

THESE MODELS ARE RETRAINED FROM SCRATCH. This is an ablation and no number here may be quoted as deployed performance; the deployed ensemble saw every organism and cannot be evaluated on any of them.

Scenario A collapses from 11.4% to 0.9% under a taxonomy-only shift. That reproduces the PET signature in a controlled setting and retires several PET-specific explanations at once: the collapse needs neither 10% sequence identity, nor the assay change, nor ESMFold structures.

RESIDUAL CONFOUND: the prokaryote arm trained on 938 fewer records (about 5%) than the control, because training proteins sharing a PIDE20 cluster with the holdout had to be dropped to prevent leakage.

### S4. Validation of the explanation axis

*The coherence axis is a construct with no ancestor in the earlier antimicrobial or transition-metal work, so it was tested rather than assumed. The established prediction-explanation direction criterion was evaluated on this property and does not transfer, which is what justifies adopting a protein-specific explanation axis. Because the axis reads a single checkpoint while the prediction reads five, its sensitivity to that choice was measured rather than argued.*

**Table S6. Sensitivity of the trust taxonomy to the choice of attribution member, on the held-out test set. Prediction is a five-model ensemble but occlusion runs on a single checkpoint, so the whole taxonomy was recomputed once per member over the identical 2,000-protein calibration cohort and the identical 2,004-protein test set. The five deployed members are listed with the checkpoint that ranked highest on held-out test but is not an ensemble member; the latter was selected using the test set and so is reported for completeness only, and no claim rests on it.**

| **attribution member** | **CV Spearman** | **in ensemble** | **coverage A/B/C/D (%)** | **MAE A/B/C/D (C)** | **A+B vs C+D** | **label agreement (%)** | **band crossings** | **residue rank rho** |
| --- | --- | --- | --- | --- | --- | --- | --- | --- |
| fold_1_repeat_0 (published) | 0.6714 | yes | 8.5 / 20.6 / 23.3 / 47.7 | 2.95 / 3.67 / 4.07 / 4.48 | 1.255x | reference | 0 | reference |
| fold_3_repeat_0 | 0.6708 | yes | 10.6 / 18.5 / 22.6 / 48.3 | 3.36 / 3.52 / 3.84 / 4.58 | 1.255x | 73.0 | 0 | 0.198 |
| fold_14_repeat_2 | 0.6707 | yes | 9.8 / 19.3 / 23.9 / 47.0 | 3.11 / 3.64 / 3.68 / 4.69 | 1.255x | 73.3 | 0 | 0.260 |
| fold_5_repeat_0 | 0.6579 | yes | 11.2 / 17.9 / 22.9 / 48.1 | 3.15 / 3.66 / 3.92 / 4.55 | 1.255x | 69.7 | 0 | 0.205 |
| fold_2_repeat_0 | 0.6448 | yes | 10.0 / 19.1 / 24.0 / 46.9 | 3.20 / 3.60 / 3.80 / 4.63 | 1.255x | 75.6 | 0 | 0.173 |
| fold_8_repeat_1 | 0.657 | no | 9.2 / 19.9 / 23.7 / 47.2 | 3.02 / 3.66 / 3.74 / 4.65 | 1.255x | 75.2 | 0 | 0.253 |

The ordering of error across scenarios replicates in every arm without exception, and Scenario A carries the lowest error in all six. Each arm recalibrates its own coherence cutoff on the shared pool, because coherence distributions shift between checkpoints and reusing a single frozen cutoff would turn a scale difference into an apparent coverage difference.

Band crossings count proteins that move between the trusted band and the untrusted band. The column is zero throughout, which follows from construction rather than from luck: the consensus axis reads the same five deployed models whatever performs the occlusion, so it fixes both band totals, and every disagreement between arms is a reallocation of A against B or of C against D. The decision the framework supports, which is whether to act on a prediction, is therefore invariant to this choice.

Two limits belong with the table. Agreement between arms should be read against a tie-order floor of 95.1%, measured by rerunning the published member after a sorting change that affects proteins with a contact-degree tie at the core boundary. Against that floor the residual disagreement is real, so an individual protein's assignment to A rather than B is not stable across checkpoints even though the aggregate ordering is. Per-residue attributions agree at a median rank correlation of about 0.20 between members, rising to 0.33 when disjoint subsets of members are averaged before comparison; both are below the value the same criterion returns for a comparable graph network on small molecules, and per-residue attributions are accordingly reported as population-level evidence rather than as a per-residue claim about any single protein.

**Table S7. Prediction-explanation consistency (criterion 3 of the antimicrobial XAI evaluation framework), adapted to regression following the transition-metal work: a prediction above the dataset mean should carry predominantly positive attributions. Both halves of the dual criterion are computed.**

| **cohort** | **n** | **consistency mean signed** | **consistency sign fraction 70pct** | **consistency dual** | **p vs chance** |
| --- | --- | --- | --- | --- | --- |
| ALL (full dataset) | 20183 | 0.4506 | 0.0344 | 0.4506 | 9.8e-45 |
| test partition | 2004 | 0.4416 | 0.0369 | 0.4416 | 1.88e-07 |
| pool partition | 18179 | 0.4516 | 0.0342 | 0.4516 | 6.45e-39 |
| scenario A | 1799 | 0.3963 | 0.0306 | 0.3963 | 1.35e-18 |
| scenario B | 4193 | 0.3549 | 0.0143 | 0.3549 | 8.690000000000001e-80 |
| scenario C | 4325 | 0.5147 | 0.0442 | 0.5147 | 0.0554 |
| scenario D | 9866 | 0.4731 | 0.0394 | 0.4731 | 9.98e-08 |
| \|pred-ref\| Q1 | 5046 | 0.4752 | 0.0329 | 0.4752 | 0.000455 |
| \|pred-ref\| Q2 | 5045 | 0.462 | 0.0339 | 0.462 | 7.42e-08 |
| \|pred-ref\| Q3 | 5046 | 0.4705 | 0.0602 | 0.4705 | 2.89e-05 |
| \|pred-ref\| Q4 | 5046 | 0.3948 | 0.0107 | 0.3948 | 8.29e-51 |

THE CRITERION IS NOT SATISFIED: consistency is 0.4506 over the full dataset, below the 0.50 chance line. That is the useful outcome, because it shows the established direction criterion does not transfer to a global emergent property such as Tm, which is what justifies adopting a protein-specific explanation axis rather than importing the molecular one.

MECHANISM: 61.9% of proteins carry a positive mean signed attribution, a systematic occlusion bias independent of Tm, which mechanically caps consistency below 50% for the below-mean half. Stratifying by distance from the pool mean shows consistency is WORST furthest from the mean (0.395) rather than nearest it, so this is not an artifact of proteins sitting on the boundary.

READ THE PER-SCENARIO ROWS WITH CARE. Scenario A scores 0.396 against Scenario D's 0.473, which could be misread as trusted predictions having worse explanations. The criterion is below chance in EVERY stratum, so it is not a yardstick Scenario A fails but an instrument carrying no signal for this property.

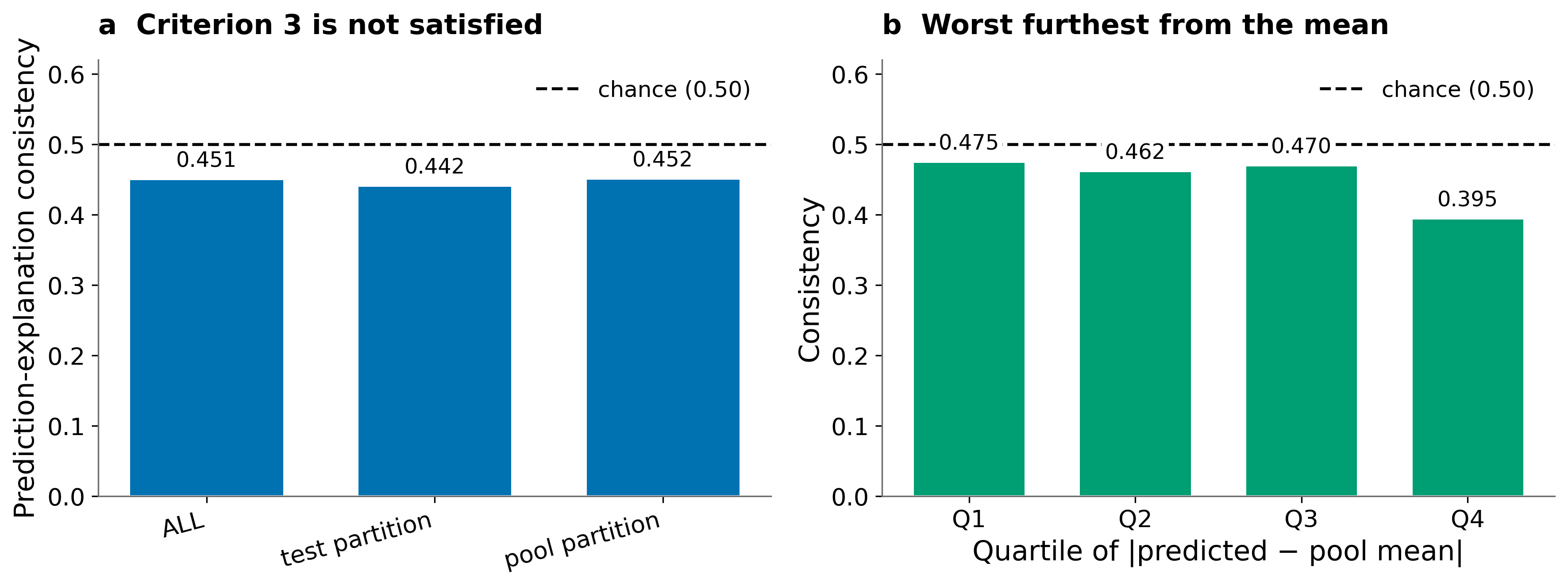

**Figure S1.** Evaluation of the prediction-explanation direction criterion on melting temperature. (a) Per-protein consistency by partition against the 0.50 chance level. (b) Consistency stratified by distance from the dataset mean; it is worst furthest from the mean rather than nearest it, so the criterion fails everywhere rather than at a decision boundary.

### S5. Model comparison and the industrial decision

*The paired comparison scores both models on identical proteins, resampling them on a shared bootstrap index, which is the correct test when two models are evaluated on one shared set. The industrial decision table is a demonstration at a stated n and supports no significance claim.*

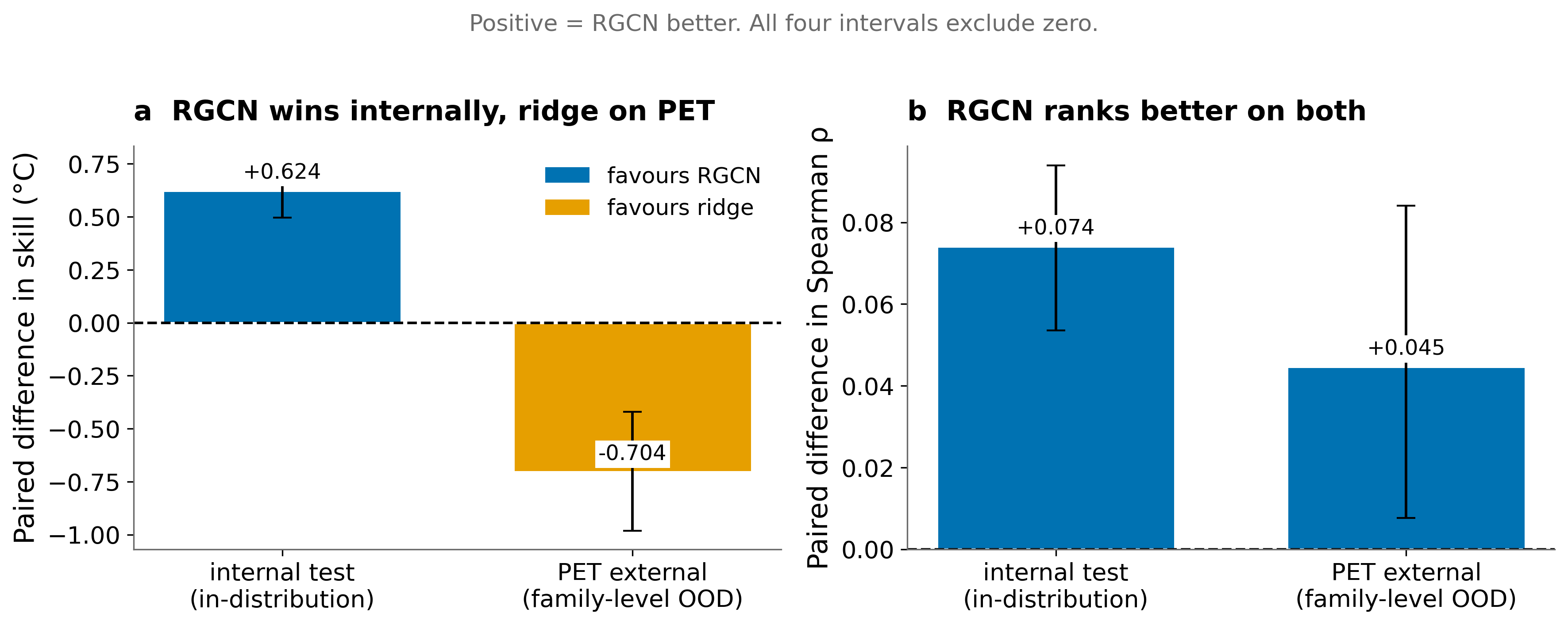

**Figure S2.** Paired comparison of the relational graph network against a mean-pooled embedding ridge on the identical proteins, resampling both models on a shared bootstrap index. (a) Paired difference in error saved relative to a constant predictor. (b) Paired difference in Spearman correlation. Positive values favour the graph model; all four intervals exclude zero.

**Table S8. The constrained industrial decision on PET: enzymes that are expressed AND above 60 C AND active against PET. Both arms are restricted to the 190 enzymes with all three endpoints measured, giving an honest base rate of 26/190 = 0.137.**

| **budget** | **arm** | **pool size** | **k effective** | **hits** | **precision** | **enrichment factor** | **p hypergeom** | **pool exhausted** |
| --- | --- | --- | --- | --- | --- | --- | --- | --- |
| 5 | score_only | 190.0 | 5.0 | 1.0 | 0.2 | 1.46 | 0.525 | False |
| 5 | trust_strict_AB | 4.0 | 4.0 | 0.0 | 0.0 | 0.0 | 1.0 | True |
| 5 | trust_strict_AB_in_domain | 4.0 | 4.0 | 0.0 | 0.0 | 0.0 | 1.0 | True |
| 5 | trust_rankmapped_AB | 54.0 | 5.0 | 0.0 | 0.0 | 0.0 | 1.0 | False |
| 5 | trust_rankmapped_AB_in_domain | 12.0 | 5.0 | 0.0 | 0.0 | 0.0 | 1.0 | False |
| 5 | random |  |  |  | 0.132 | 0.97 |  |  |
| 10 | score_only | 190.0 | 10.0 | 2.0 | 0.2 | 1.46 | 0.409 | False |
| 10 | trust_strict_AB | 4.0 | 4.0 | 0.0 | 0.0 | 0.0 | 1.0 | True |
| 10 | trust_strict_AB_in_domain | 4.0 | 4.0 | 0.0 | 0.0 | 0.0 | 1.0 | True |
| 10 | trust_rankmapped_AB | 54.0 | 10.0 | 1.0 | 0.1 | 0.73 | 0.779 | False |
| 10 | trust_rankmapped_AB_in_domain | 12.0 | 10.0 | 0.0 | 0.0 | 0.0 | 1.0 | False |
| 10 | random |  |  |  | 0.133 | 0.97 |  |  |
| 20 | score_only | 190.0 | 20.0 | 4.0 | 0.2 | 1.46 | 0.284 | False |
| 20 | trust_strict_AB | 4.0 | 4.0 | 0.0 | 0.0 | 0.0 | 1.0 | True |
| 20 | trust_strict_AB_in_domain | 4.0 | 4.0 | 0.0 | 0.0 | 0.0 | 1.0 | True |
| 20 | trust_rankmapped_AB | 54.0 | 20.0 | 3.0 | 0.15 | 1.1 | 0.537 | False |
| 20 | trust_rankmapped_AB_in_domain | 12.0 | 12.0 | 0.0 | 0.0 | 0.0 | 1.0 | True |
| 20 | random |  |  |  | 0.136 | 0.99 |  |  |

The previously reported base rate of 0.0547 was DILUTED: it divided by all 475 enzymes, of which 285 could never be counted as hits because at least one endpoint was unmeasured. That figure must not be quoted.

NO SIGNIFICANCE CLAIM IS AVAILABLE AT ANY BUDGET. This is a demonstration at a stated n, not a test. An arm marked pool_exhausted nominated fewer candidates than the budget, so its hit rate is computed over fewer enzymes and is not comparable to the others; the strict trust-aware arm nominates only 4 enzymes out of 190, which is the framework declining to recommend rather than a defective table.

Two readings survive. Ranking by predicted Tm alone outperforms every trust-filtered arm on this family-level out-of-distribution set, which is what the conditional deployment rule predicts. And adding the applicability-domain filter cuts the rank-mapped pool from 53 to 12 and its hits from 3 to 0, independently reproducing the measured inversion of that axis.
